# T cell development from expanded hematopoietic progenitors reveals progression control by *Lmo2*, *Erg*, *Spi1*, *Hoxa9*, and *Meis1*

**DOI:** 10.1101/2025.04.22.649893

**Authors:** Boyoung Shin, Samantha J. Chang, Brendan W. MacNabb, Tom Sidwell, Brian A. Williams, Ellen V. Rothenberg

**Affiliations:** Division of Biology & Biological Engineering, California Institute of Technology, Pasadena, CA 91125; Single-cell Profiling and Engineering Center, Beckman Institute, California Institute of Technology, Pasadena, CA 91125

**Keywords:** Expanded hematopoietic stem and progenitor cells, early T cell program, chromatin accessibility, single-cell transcriptome analysis, Lmo2, Hoxa9, Meis1, Flt3L, germline TCRβ transcription

## Abstract

To gain access to the earliest stages of T cell development, we adapted a serum-free culture system that expands hematopoietic stem and progenitor-like cells. These expanded cells efficiently undergo normal T-cell differentiation *in vivo* and *in vitro*, verified by early gene expression trajectories from single-cell RNA sequencing, though their absolute differentiation speed is slower than that of fresh progenitors and can be modulated with cytokine priming. Leveraging this expansion system to observe the first T-lineage events, we revealed that initial Notch activation immediately induces chromatin opening and transcriptional activation of the TCR-Cβ locus. Additionally, acute CRISPR knockouts confirmed T-lineage entry requirements for *Ikzf1*, *Hes1*, *Gabpa*, and *Myb* while revealing that *Lmo2*, *Erg, Spi1, Hoxa9*, and *Meis1* retard developmental progression with differing effects on proliferation. Endogenous expression of the stem, progenitor, and leukemia-associated factor Lmo2 markedly restrains initiation of the T cell program, with *Lmo2* knockout greatly accelerating germline TCRβ locus transcription and expression of *Tcf7, Gata3,* Runx family, and E protein genes and their targets.

## INTRODUCTION

The earliest molecular events that cause hematopoietic progenitors to enter the T cell pathway have remained poorly understood. T cells share a common progenitor with other blood cells: the bone marrow hematopoietic stem cell (HSC). However, T cell development occurs in a physically separated organ, the thymus, where only a tiny number of hematopoietic precursors initiates T cell development per day, resulting in obvious developmental discontinuity from other hematopoietic lineage cells. Previous studies have characterized diverse progenitor cell types in bone marrow that include candidate prethymic precursors as well as the different developmental stages of intrathymic T-progenitor cells as they start the T cell program (*1–4*). The cells seeding the thymus (*5–8*) go through both stepwise and continuous developmental changes as they proliferate while receiving strong Notch signaling and cytokine/chemokine cues from the thymic microenvironment (*9–13*), starting from the initial CD4 and CD8 double negative (DN) stages (DN1, DN2a, DN2b, DN3a, DN3b, and DN4, Fig. 1A). During the pro-T cell stages, DN1-DN3a, progenitor cells undergo substantial developmental and epigenomic conversion from the uncommitted, multipotent state (DN1 and DN2a) to a T-lineage committed state (DN2b and onward), even before detectable cell-surface T cell receptor (TCR) expression. Yet, while single-cell transcriptome data from prethymic and intrathymic cells suggest possible connections with particular bone marrow precursors (*14–17*), the initial events that occur in immigrating cells upon thymic entry remain unclear.

**Figure 1.**
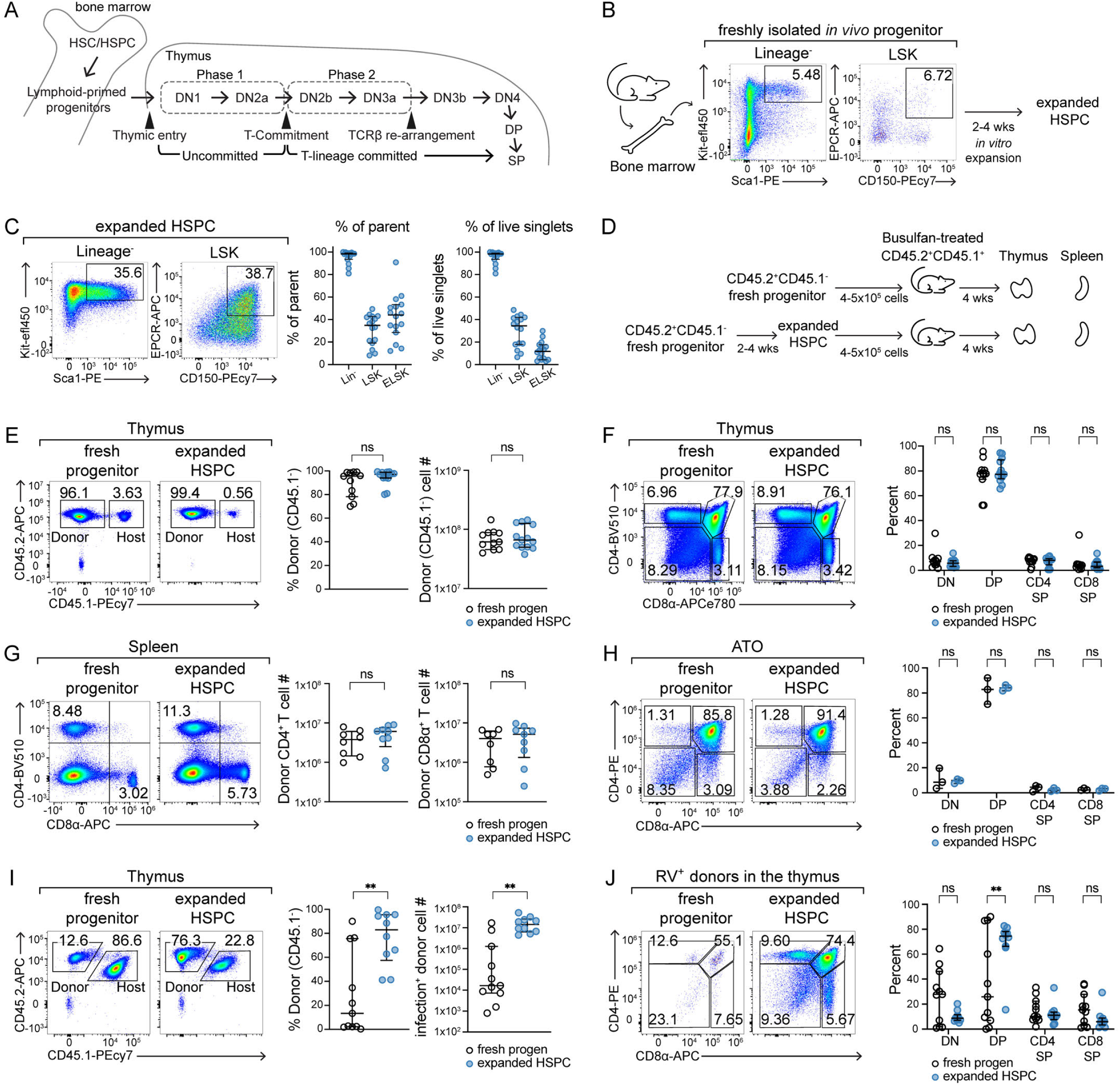
Expanded HSPCs are competent to generate T cells *in vivo* and *in vitro*. **(A)** Graphical illustration of T cell development. Thymic T cell development from double negative (DN), double positive (DP), to single positive (SP) stages and major events are shown (e.g., thymic entry). The uncommitted multipotent progenitor stages (“Phase 1”) and T-lineage committed stages before TCRβ rearrangement (“Phase 2”) are marked. HSC: Hematopoietic stem cells, HSPC: hematopoietic stem and progenitor cells. **(B)** Graphical diagram with representative flow cytometry plots show HSPC isolation and expansion workflow. **(C)** Representative flow cytometry plots show Kit, Sca1, EPCR, and CD150 expression profiles of expanded HSPCs after 2-4 weeks of expansion. Graphs summarize median values of frequencies of lineage^-^ (Lin^-^), Lin^-^ Sca1^+^Kit^+^ (LSK), and EPCR^+^CD150^+^LSK cells with interquartile ranges from 16 independent experiments. **(D)** A schematic diagram shows bone marrow chimera reconstitution and analysis strategies. **(E-G)** Freshly isolated Lin^-^ progenitors or Lin^-^ expanded HSPCs from CD45.1^-^ CD45.2^+^ mice were transferred into busulfan-conditioned CD45.1^+^CD45.2^+^ recipients. Donor-derived cell profiles in the thymus and the spleen were analyzed after 4 weeks of reconstitution. 2-3 independent experiments (n=3-5 mice for each group per experiment, a total of 11-13 mice). Median values and interquartile ranges are shown in the graphs. **(E)** Reconstitution efficiencies of fresh progenitors and expanded HSPCs were assessed by analyzing the thymus of the recipients using flow cytometry. The frequency of CD45.1^-^CD45.2^+^ cells was determined by gating on live singlet lymphocytes. Statistical comparisons by unpaired t-test. ns = not significant. **(F, G)** CD4 and CD8α expression patterns of donor cells in the thymus (F) and the spleen (G) of recipient mice were measured using flow cytometry. **(F)** Percentages of indicated populations in the thymus. Comparisons by two-way ANOVA with Šidák correction for multiple comparisons. ns = not significant. **(G)** Representative flow cytometry plots display splenic CD4 and CD8 T cell frequencies. Graphs summarize donor-derived splenic CD4 and CD8 T cell numbers. Comparisons by unpaired t-test. ns = not significant. **(H)** *In vitro* T-development of fresh progenitors vs. expanded HSPCs was examined using an artificial thymic organoid (ATO) system. Percent of DN, DP, CD4 SP, and CD8 SP were determined using flow cytometry after 4 weeks of ATO culture. The graph summarizes 3 independent experiments (each dot = mean of 1-5 technical replicates). Two-way ANOVA with Šidák correction for multiple comparisons. ns = not significant. **(I, J)** Evaluating T-developmental potentials of retrovirally engineered donor cells. Fresh progenitors or expanded HSPCs were infected with retroviral vectors, and 1×10^5^ of infected Lin^-^ cells were sorted and transferred into congenically marked busulfan-treated recipients. After 5-6 weeks post-transfer, donor cells in the thymus were analyzed. Median values and interquartile ranges are shown from 3 independent experiments with 3-4 mice per group, a total of 10-11 mice. **(I)** Representative flow cytometry plots display the retrovirally transduced donor cell reconstitution rates in the thymus. The graphs summarize the donor cell frequency and infection^+^ donor cell number in the thymus. Statistical significance was assessed by unpaired t-test with Welch’s correction for unequal standard deviation. ** = *p-*value < 0.01, ns = not significant. **(J)** Thymic CD4 and CD8α expression are shown with frequencies overlaid. The graph displays the percent of different thymic populations with median and interquartile ranges. Two-way ANOVA with Šidák correction for multiple comparisons. ** = *p-*value < 0.01, ns = not significant.

Recent single-cell transcriptome data have uncovered intriguing heterogeneity among cells in the true early T-cell progenitor (ETP) subset of DN1 thymocytes, but those progressing into the T-cell program converge on a common developmental trajectory (*16, 18, 19*). This convergence depends on the first molecular events in development, partially explained by the Notch signaling that thymic-seeding progenitors receive, which is vital for both the competence to enter the T-cell program (*20*) and for developmental progression (rev. in (*21*)). However, chromatin accessibility patterns suggest that pro-T cells remain extremely close to prethymic lymphomyeloid precursors despite this Notch signaling, not only in the ETP stage, but up to commitment in DN2b stage (*22, 23*). Thus, the initiation of T cell development proper remains incompletely understood. This developmental transition has been challenging to study at a molecular level due to the scarcity of relevant bone marrow progenitor cells and pro-T cells, heterogeneity of progenitor populations, dilution of early-stage cells by proliferation as cells progress in development, and the speed with which cells exit the earliest stages of T cell development when studied *in vitro*.

Here, we show that with slight modification, a recently reported method for expanding multipotent hematopoietic progenitors *in vitro* generates large numbers of enriched T-lineage competent precursors that are highly tractable for genetic manipulation. First, we comprehensively characterize T cell development from these progenitors using flow cytometry, population RNA-seq, single-cell RNA-seq and ATAC-seq. Next, we use this system to reveal the earliest gene expression and chromatin accessibility changes as T-cell development starts. We demonstrate an accelerating role for Flt3L stimulation. Then, using this system, we show that the kinetic impacts of multiple progenitor-factor regulators in the earliest stages of T-lineage entry can be measured with highly improved sensitivity, and effects on differentiation in real time can be clearly distinguished from selective effects on population expansion. This system confirms effects of PU.1 and Erg on the kinetics of T-cell developmental initiation, and reveals novel early roles for Hoxa9, Meis1, and especially Lmo2.

## RESULTS

### The HSC expansion method supports the proliferation of bone marrow progenitors while preserving T cell potentials

We first asked whether a recent technical innovation affording ∼230-to-800-fold expansion of HSCs (*24, 25*) could overcome existing cell number limitations by providing an appropriate, accessible platform for monitoring the transition from the pre-thymic bone marrow progenitor state to the early thymic pro-T cell state (Fig. 1A).

We isolated bone marrow progenitors that were lineage marker negative (“Lin^-^”, defined by TCRβ^-^ TCRγδ^-^ CD19^-^ CD11b^-^ CD11c^-^ Ly6G/C^-^ NK1.1^-^ CD49b^-^ and Ter119^-^), but Sca1^+^Kit^+^(“LSK”), and also positive for CD150 and endothelial protein C receptor (EPCR or PROCR/CD201), highly expressed surface molecules in HSCs (*26–31*). These progenitors were expanded under serum-free conditions supplemented with thrombopoietin (TPO, 100 ng/mL), stem cell factor (SCF, 10 ng/mL), and polyvinyl alcohol (PVA), as described (*24, 25*) (Fig. 1B).

After 2-4 weeks of expansion, most of the expanded hematopoietic stem and progenitor cells (HSPCs) were still negative for expression of any mature hematopoietic lineage markers (Lin^-^; median 98.25%); ∼20-50% of these Lin^-^ cells displayed the Lin^-^ Sca-1^+^ Kit^+^ “LSK” phenotype (median 34.75%), and ∼30-50% of LSK cells expressed both CD150 and EPCR (“ELSK”; median 44.1%) (Fig. 1C). Previous studies had reported that expanded HSPCs with EPCR^+^(also *Fgd5*^+^) LSK phenotypes closely mimicked the gene expression signatures with those of freshly isolated HSCs, and had superior repopulation capacity upon *in vivo* transfer (*32, 33*). Therefore, we further isolated LSK or ELSK cells again from the expansion cultures to serve as the expanded HSPC inputs for most of our experiments.

To assess whether expanded HSPCs truly retain T cell potential after the expansion period, we generated bone marrow chimeras by transferring expanded HSPCs or freshly isolated bone marrow progenitors into busulfan-treated, congenically distinct recipients (4-5 x 10^5^ cells per mouse) (Fig. 1D). At 4 weeks post-transfer, both types of donor cells efficiently reconstituted the recipients’ thymus (Fig. 1E ∼95.5-97% reconstitution rate) and generated essentially equal frequencies of double negative (DN), double positive (DP), CD4- and CD8-single positive (SP) populations (Fig. 1F). Moreover, both expanded HSPCs and freshly isolated progenitors not only produced TCRβ^+^ and TCRγδ^+^ T cells in the thymus (Fig. S1A), but also started to repopulate the T cell compartment of the spleen with equal frequencies (Fig. 1G, S1B). To test whether expanded HSPCs could also generate mature-phenotype T cells during differentiation *in vitro* for experimental tracking and monitoring, we used an artificial thymic organoid (ATO) 3D culture system (*34, 35*). Consistent with *in vivo* results, the ATOs formed by expanded HSPCs generated DN, DP, CD4 SP, and CD8 SP cells, including TCRβ^+^ and TCRγδ^+^ T cells, at least as well as those formed by freshly isolated progenitor cells (Fig. 1H, S1C). Thus, *in vitro* expanded HSPCs could efficiently develop into mature T cells both *in vivo* and *in vitro*.

Since expanded HSPCs retained T cell potential even after multiple rounds of cell divisions, we next asked whether retroviral (RV) vectors could be utilized in expanded HSPCs to engineer or perturb gene expression in the progenitor cells that give rise to T cells in vivo. In freshly isolated populations, the cells susceptible to retroviral transduction are often poor at reconstituting recipient animals, likely because transduction requires cycling cells and the best engrafting progenitors in the adult bone marrow are not in cell cycle. To test how cultured progenitors respond to RV-transduction, expanded HSPCs and freshly isolated Lin^-^ progenitors were infected with empty-vector control retrovirus overnight. Next, cells positive for the RV infection marker were sorted and transferred into busulfan-treated, CD45.1^+^CD45.2^+^ congenically marked recipients (10^5^ infection-positive Lin^-^ cells per mouse) (Fig. S1D). After 5-6 weeks post-transfer, as expected, freshly isolated progenitors after infection were inefficient in repopulating the host thymus (median donor-type CD45.1^-^CD45.2^+^ percentage: 13.40 %), with a poor DP:DN cell ratio, suggesting improper T cell development (Fig. 1I, J). In contrast, expanded HSPCs after infection successfully reconstituted the host thymus (median donor-type CD45.1^-^ CD45. 2^+^ percentage: 82.95 %) and efficiently generated DN, DP, CD4 SP, and CD8 SPs (Fig. 1I, J).

Furthermore, most of the cells that did differentiate from infection-positive, freshly isolated donor cells had silenced expression of the retroviral DNA (7 out of 11 recipient mice showed <10% infection marker positivity). In contrast, RV-transduced expanded HSPCs maintained infection marker expression in all tested recipient animals (Fig. S1D). Thus, this PVA-based HSPC expansion method provides a new opportunity to investigate T cell development following retroviral vector-based progenitor engineering.

### Expanded HSPC descendants pass through early T-lineage developmental stages with slowed kinetics

We tested whether expanded HSPCs and fresh progenitors initiate similar T cell developmental programs by tracking the behavior of their respective progeny after exposure to early T-development conditions *in vitro*. Briefly, each group of progenitors was co-cultured with OP9 stromal cells expressing the Notch ligand Delta-like 1 (Dll1), in medium supplemented with Flt3-ligand (Flt3L) and IL-7, a well-established *in vitro* system closely recapitulating early stages of intrathymic T cell development (*36*)(direct transcriptome comparison with *in vivo* populations in (*37*)). In these T lineage-inducing conditions, expanded HSPC grew with approximately 1/5 cloning efficiency (1/2.2-1/8.5 in three experiments; Fig. S1E). After 4-10 days of co-culture, early T-developmental progression state was followed using the surface markers Kit, CD25, and Thy1/CD90 expression to trace the cells from ETP (Kit^+^ DN1 cells) through DN3 stage (CD25^+^ Thy1^+^ Kit^low^), and using the *Bcl11b*-mCitrine reporter (a non-disruptive, bicistronic knock-in allele) to highlight the transition to T-cell lineage commitment between DN2a and DN2b as previously described (*38*)(Fig. 2A).

**Figure 2.**
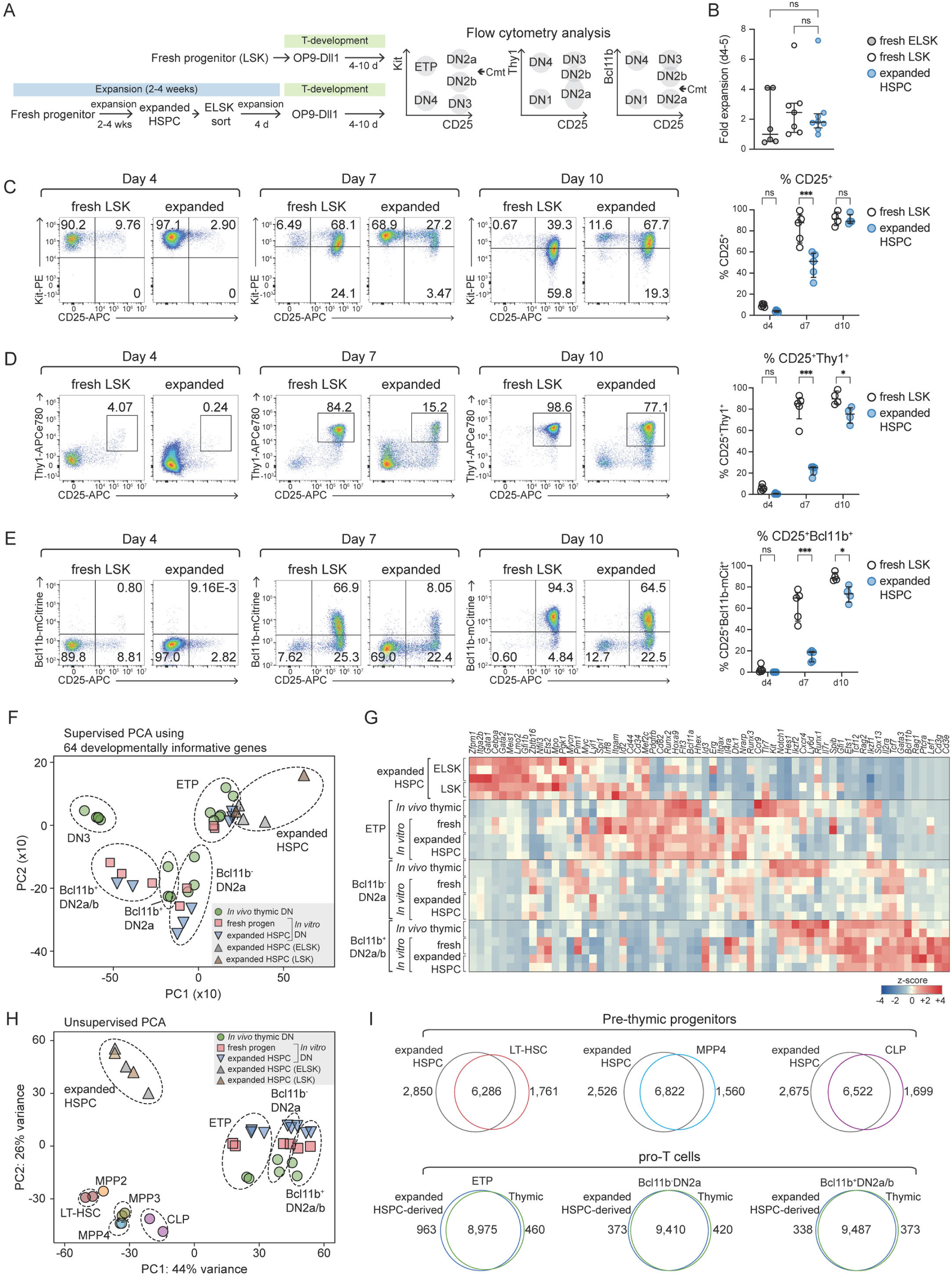
The progeny of expanded HSPCs go through early T-development stages with slow kinetics, yet they converge gene expression profiles with normal T-development programs. **(A)** The graphical scheme shows an *in vitro* OP9-Dll1 co-culture strategy to recapitulate early T cell development from fresh progenitors or expanded HSPCs. LSK cells from freshly isolated progenitors or CD150^+^EPCR^+^LSK cells from HSPC expansion were sorted and co-cultured with OP9-Dll1 for 4-10 days. T-developmental progression was determined using Kit, CD25, Thy1, and *Bcl11b*-mCitrine reporter expressions. **(B)** Net population growth of fresh bone marrow ELSK or LSK progeny and expanded HSPC progeny after 4-5 days of OP9-Dll1 co-culture. Comparisons by Kruskal-Wallis test. ns = not significant. **(C-E)** Pro-T cells from OP9-Dll1 co-culture were analyzed on days 4, 7, and 10 using flow cytometry. Representative flow plots for each timepoint are shown with frequencies. The frequencies of CD25^+^, CD25^+^Thy1^+^, and CD25^+^*Bcl11b*-mCitrine^+^ cells from 5 independent experiments were summarized with median and interquartile ranges. Statistical analysis by two-way ANOVA with Šidák correction for multiple comparisons. *** = *p-*value < 0.001, * = *p-*value < 0.05, ns = not significant. **(F-I)** Transcriptional profiles of expanded HSPCs and their progeny DN populations were examined using bulk RNA-seq. Expanded HSPCs were sorted for CD150^+^EPCR^+^LSK or LSK phenotypes (noted as ELSK or LSK) to measure transcriptional profiles. To obtain pro-T cells from expanded HSPCs, expanded HSPCs (LSK) were subjected to OP9-Dll1 co-culture, and ETP (Kit^+^CD25^-^Bcl11b-mCitrine^-^), Bcl11b^-^ DN2a (Kit^high^CD25^+^Bcl11b-mCitrine^-^) and Bcl11b^+^ DN2a/b (Kit^high/int^CD25^+^Bcl11b-mCitrine^+^) cells were sorted on day 5 or day 9 and subjected to bulk RNA-seq. Gene expression programs of *in vivo* progenitors from the bone marrow (*22*), *in vivo* thymic progenitors (*18*), and *in vitro* DNs from fresh progenitors (*39*) were obtained from previously published datasets. **(F, G)** Supervised principal component analysis (PCA) using 64 T-developmental trajectory genes shows the positions of expanded HSPCs and their progeny on the known developmental pathway. The heatmap displays expression levels of the 64 genes in indicated progenitor populations (G). **(H)** Unsupervised PCA illustrates the variances across *in vivo* progenitor populations in the bone marrow, the thymus, and *in vitro* pro-T cells on the PC1 and PC2 axes. **(I)** Area-proportional Venn diagrams show the number of differentially expressed genes (crescent area) and not differentially expressed genes (intersect area) from indicated comparisons.

Net population growth was similar in both sets of cultures from d4 – d10 (Fig. 2B), but the cohorts derived from these two different progenitors went through ETP to DN2b stages at different rates (Fig. 2C-E). On day 4, progeny of both freshly isolated LSKs and expanded HSPCs predominantly kept the ETP surface phenotype. By d10, some progeny of both input progenitors developed to DN2 and DN3 stage pro-T cells, successfully upregulating CD25 and *Bcl11b* (Fig. 2C-E). However, induction of CD25, Thy1, and *Bcl11b*-mCitrine reporter was delayed by several days in expanded HSPC-derived pro-T cells as compared to pro-T cell progeny of freshly derived precursors.

Furthermore, despite their ability to progress to mature T cells in the ATO system (Fig. 1H, Fig. S1B), progeny of expanded HSPCs spent a more prolonged period in the ETP stage than progeny of fresh precursors, and more slowly progressed to the DN2 and DN3 stages in that system as well (Fig. S2A, B).

### Transcriptional profiles of expanded HSPCs undergoing T cell development program converge with those of *in vivo* thymocytes

These results raised the question of whether there are abnormalities in the developmental trajectory of expanded HSPCs that would prevent them from being a useful model for probing the initiation of T cell development. We therefore compared gene expression profiles in the input expanded HSPCs and each resulting pro-T cell population after OP9-Dll1 co-culture (ETP, Bcl11b^-^DN2a, and Bcl11b^+^DN2a/DN2b), using bulk RNA-sequencing (RNA-seq) of sorted populations. These RNA expression results were compared to published reference datasets for fresh thymocytes and for cells differentiated *in vitro* from freshly isolated precursors (*18, 22, 37*).

We first used supervised principal component analysis (svPCA) based on 64 highly informative developmental regulatory genes from a reference single-cell RNA-seq (scRNA-seq) analysis of adult *in vivo* thymocytes (*18*). This approach previously allowed us to capture both normal and abnormal developmental trajectories across ETP to DN3 stages in a fixed frame of PC1 and PC2 by projecting onto defined PC loadings (*39, 40*). Here, despite the observed temporal differences, pro-T cells generated by expanded HSPCs (triangle symbols) mapped close to the corresponding control populations of *in vivo* thymocytes (square symbols) and their *in vitro* counterparts produced by freshly isolated progenitors (circle symbols) (Fig. 2F). Thus, expanded HSPC-derived DN populations robustly induced genes coding for transcription factors necessary for specifying T cell fate (e.g., *Tcf7*, *Gata3*, *Bcl11b*, *Tcf12*, and *Ets1*) and DN2/DN3 signature genes (e.g., *Il2ra*, *Cd3g*, *Cd3e*, *Ptcra*, and *Rag1*)(Fig. 2G). They also effectively downregulated stem and progenitor-associated genes (e.g., *Erg*, *Mef2c*, *Hoxa9*, *Cd34*, *Flt3*, *Spi1*, *Kit*, *Hhex*, *Bcl11a*, and *Cd44*) as they developed from ETP to DN2b stages (Fig. 2G).

To explore the source of the observed kinetic differences, we next compared gene expression programs of expanded HSPCs with those of *in vivo* progenitors from the bone marrow and the thymus (*18, 22*) using unsupervised PCA (Fig. 2H). Notably, this revealed that expanded HSPCs were not equivalent to purified bone marrow progenitors reported by the ImmGen consortium (*22*), as mapped in PCA space (Fig. 2H). The differences were most captured by the PC2 values, which were strongly driven by the higher expression of megakaryocyte, myeloid, and progenitor genes such as *Pf4*, *Jund*, *Cebpb*, and *Itga2b* in expanded HSPCs. (Supplementary Table 1). However, after engaging with the Notch-signaling environment, pro-T cell progeny of expanded HSPCs readily clustered with *in vivo-*derived thymic pro-T cells (Fig. 2H). This gene-expression convergence upon exposure to early T-development conditions was supported by differential gene expression tallies. Expanded HSPCs vs. *in vivo* bone marrow progenitor comparisons, representing pre-thymic stages, found over 4,000 differentially expressed genes (DEGs)(Fig. 2I). In contrast, comparisons of pro-T cells derived from expanded progenitors in OP9-Dll1 cocultures against *in vivo* pro-T cells found only ∼1,400 DEGs in ETP stage and only ∼700 DEGs in DN2a/DN2b stages (Fig. 2I, Supplementary Table 2). The most prominent residual differences at these pro-T cell stages included higher expression of genes associated with glycolysis, oxidative phosphorylation, mTORC1, cell cycle (Myc targets), and fetal vs. adult thymic progenitor differences (Fig. S2C, D) in the expanded HSPC-derived *in vitro* pro-T cells as compared to the *in vivo* thymic pro-T cells. Thus, expanded HSPCs properly initiated the early T-development programs in response to strong Notch and cytokine signaling, generating transcriptional programs that closely resembled those of normal *in vivo* thymic progenitor cells at each DN stage, but with elevated activity of metabolism and cell cycle genes.

### Single-cell transcriptomes of progeny of expanded HSPCs and freshly isolated bone marrow HSPCs converge to the same early T-development trajectory

We next investigated whether the specific differentiation pathways leading to convergent pro-T states from expanded HSPC progenitors were consistent with those from fresh progenitors, and thus relevant for studying T-lineage initiating events. We used single-cell RNA sequencing (scRNA-seq) to track these specific differentiation pathways, comparing freshly isolated ELSK (CD150^+^EPCR^+^LSK) progenitors from the bone marrow with expanded HSPCs sorted on LSK or ELSK markers. Besides input samples, progeny of both input types in early T-cell development cultures were harvested at two different culture timepoints before sorting for scRNA-seq (Fig. 3A)(see Supplemental Methods for analysis details). Cultures were seeded so that all samples in an experiment could be harvested on the same day, barcoded, and processed in the same 10X chip to minimize batch effects. However, we integrated results of two independent experiments in the analysis and also integrated results of two independent previously published *in vitro* pro-T cell scRNA-seq datasets from fresh progenitors, to provide an additional layer of anchoring points across intermediate stages (Fig. 3A).

**Figure 3.**
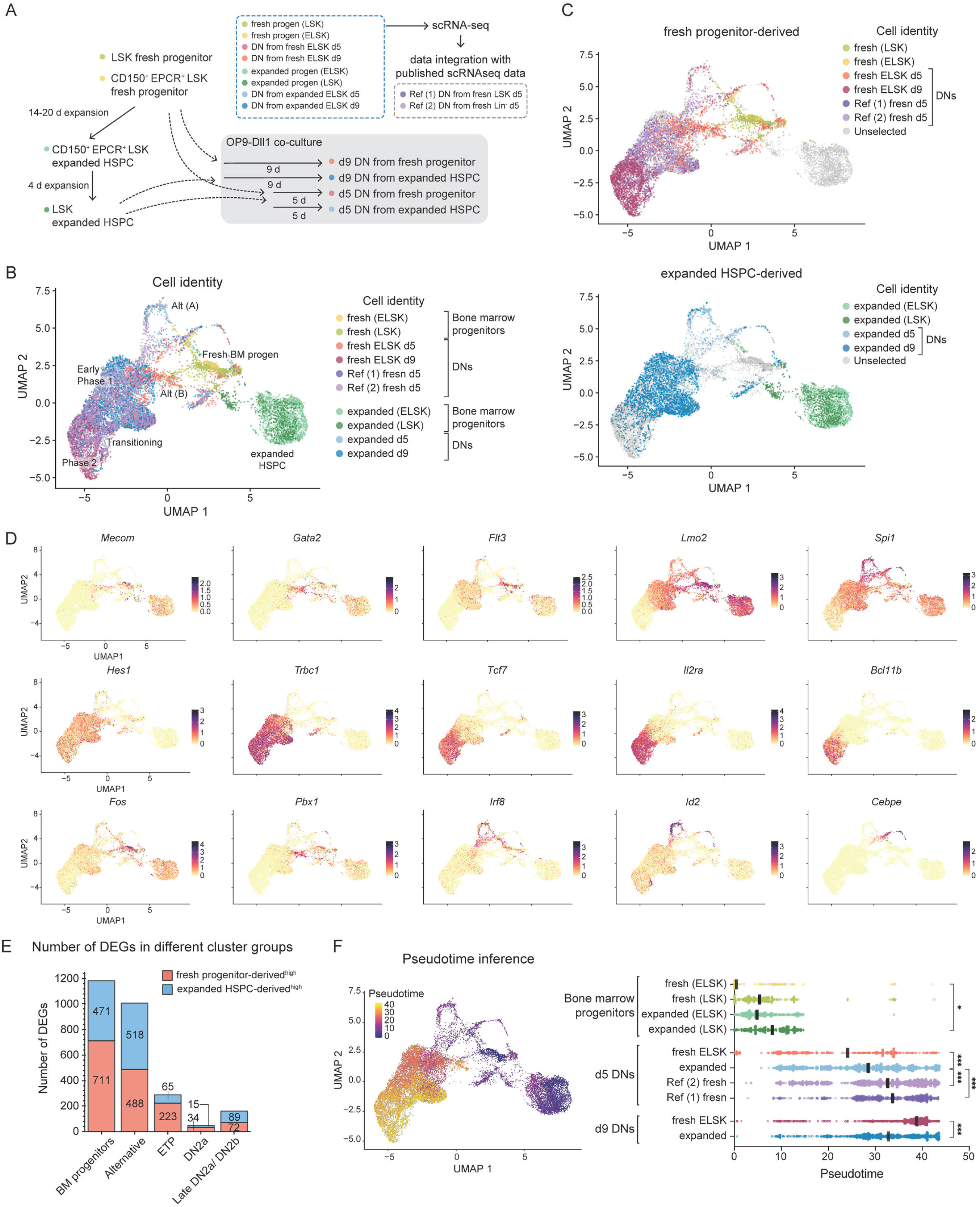
Expanded HSPCs and fresh progenitors share an early T cell development trajectory after entering the T cell program at the single-cell transcriptome level. **(A)** The schematic shows single-cell RNA-seq (scRNA-seq) experimental design. Each condition group was marked by a unique hashtag oligo (HTO) and pooled for scRNA-seq. Two independent scRNA-seq experiments were performed. Two sets of reference scRNA-seq from fresh progenitor-derived *in vitro* pro-T cells (*37, 51*) were integrated to provide additional reference points across DN1 to DN2b stages. **(B)** The identity of each condition is shown on the UMAP1-2 manifold. **(C)** Distinct types of input progenitors (fresh or expanded) are highlighted. **(D)** The expression levels of informative genes are shown by color intensity. **(E)** Stacked bar graphs depict the numbers of differentially expressed genes (DEGs) by progeny of fresh vs. expanded HSPC-descendants within the progenitors (clusters 1, 3, 7, 13, 17, 18), alternative lineages (clusters 11, 12, 14, 15, 20, 21), early Phase 1 (clusters 0, 2, 5, 6, 9), transitional (clusters 10, 16), and Phase 2 (clusters 4, 8, 19) groups. The DEGs in each group were determined by absolute Log_2_ fold-change greater > 1, and adjusted *p-*value < 0.01. **(F)** The inferred pseudotime score of each cell is shown by colormap and scatter plot. The root principal node was defined based on known LT-HSC features (*Mecom* > 3 & *Tcf15* > 1 & *Kit* > 1 & *Procr* > 1 & *Ly6a* > 1 & *Il7r* < 0.5 & *Flt3* < 0.5). Kruskal-Wallis test of multiple comparisons. *** = adj.*p-* value < 0.001.

Before encountering a Notch signaling environment, expanded HSPCs formed a relatively uniform supercluster (Fig. 3B, S3A, clusters 1, 3, and 7). As expected from the bulk RNA-seq PCA analysis, this was distinct from freshly isolated bone marrow progenitors, which comprised multiple clusters (predominantly clusters 13, 17, 18, but also included in clusters 11, 12, 14, and 15) (Fig. 3B, C1, S3A-C). However, freshly isolated progenitors and expanded HSPCs exhibited markedly similar gene expression programs under T-development conditions (days 5-9) and passed through the same pro-T cell clusters (Fig. 3C). After 5 days of OP9-Dll1 co-culture, pro-T cells that developed from both input populations occupied clusters 2, 5, and 9, representing early Phase 1 stage cells (∼ETP) on the T-development pathway (Fig. 3C; Fig. S3B, C). Cells in this population were marked by expression of *Notch1* and Notch signaling-induced genes (e.g. *Hes1*), and by robust induction of *Trbc1*, from one of the TCRβ germ-line transcripts (Fig. 3D, Fig. S3D). At day 9 after T-development induction, both populations transitioned toward the later transitional (∼DN2a), and Phase 2 (∼DN2b) clusters (Fig. 3C, D), ultimately turning on post-commitment genes such as *Bcl11b*, *Lef1*, *Dntt*, and *Cd3g/Cd3d/Cd3e* cluster genes (Fig. 3D, Fig. S3D).

In accord with the evidence for differences in developmental speed (Fig. 2), the progeny of freshly isolated ELSKs had mostly evacuated from the ETP stage (clusters 0, 2, 5, 6, and 9) and reached DN2a (clusters 10 and 16) and DN2b stages (clusters 4, 8 and 19) by day 9, whereas more of the progeny of expanded HSPCs persisted in the ETP stages (Fig. S3B). However, not only did the cells derived from freshly isolated and expanded HSPCs pass through the same UMAP clusters, but they also showed minimal differences in gene expression once they began T-development. Only 50-300 DEGs could be scored by pseudo-bulk DE analysis between progeny of different precursor types within each pro-T cell cluster group (Fig. 3E), confirming the convergence seen in the bulk RNA-seq results above. The similarity of trajectories enabled us to relate these changes to a uniform T-lineage development pseudotime scale calculated from the whole set of scRNA-seq data (Fig. 3F). Comparison of individual cell states along this pseudotime axis showed that the progeny of expanded HSPCs initially advanced at least as fast as the progeny of fresh ELSKs, but then progressed to later pseudotime values more slowly on average (Fig. 3F). Collectively, these results demonstrate that freshly isolated bone marrow progenitors and expanded HSPCs utilize a common developmental route during *in vitro* early T cell development despite their different developmental progression rates and different starting states.

### Different alternative lineage pathways for non-T progeny of expanded HSPC and freshly isolated HSPC

Where progeny of expanded HSPCs and freshly isolated HSPCs differed was in their generation of other, non-T cell types in OP9-Dll1 coculture. Both bone marrow progenitor-associated clusters and the non-T progeny that they generated upon differentiation showed the largest number of DEGs between fresh bone marrow-derived and expanded HSPC-derived samples (∼1100 and ∼1000 DEGs, respectively) (Fig. 3E). For example, expanded HSPC progeny cells that did not become T cells were preferentially associated with clusters 11, 15, and 21, which expressed high levels of myeloid/DC genes (*Irf8*, *Cebpb*, *Id2*, and *Mpo*), whereas progeny of freshly isolated progenitors predominantly occupied clusters 12, 14, and 20, which expressed high levels of megakaryocyte- and HSC-associated genes (*Gata2*, *Mecom*, *Pf4*, *Hlf*, and *Pbx1*) (Fig. 3D, S3B, D). These differences contrasted with the sharply decreased numbers of DEGs defined by scRNA-seq in pro-T cell progeny clusters (∼50 – 300 DEGs) (Fig. 3E). Thus, most variations between expanded HSPCs and freshly isolated HSPCs related to initial differences in alternative lineage biases.

### Developmental chromatin accessibility dynamics highly coordinated with entry to the early T cell program

With this foundation established, we exploited the expanded HSPC system to define how the earliest events in the T cell program changed the chromatin landscape. We determined the relationships between chromatin states in expanded HSPCs and their early T-lineage progeny, and identified the specific genomic regions that undergo the earliest chromatin accessibility changes as T-lineage differentiation begins (Fig. 4). Accordingly, we performed bulk ATAC-seq on sorted, expanded HSPCs and on sorted DN subsets derived from these expanded HSPCs. We further included ATAC-seq samples prepared under similar conditions from a newly-generated set of *in vivo* thymic DN populations (Fig. 4A-C). As an additional reference, we also included ATAC-seq data from *in vivo* bone marrow and thymic progenitors reported by the ImmGen consortium (*22*) in comparisons. In PCA analysis (please see Supplemental Methods for details), the PC1 values efficiently reflected developmental states, with the most immature LT-HSCs showing the highest PC1 value and the most progressed DN2b cells from all sample series displaying the lowest PC1 values (development from right to left). The ATAC-seq samples made by different groups using different methods showed significant technical variations, but these were confined to PC2 (Fig. 4A). Importantly, *in vivo* and expanded HSPC-derived *in vitro* pro-T cells were highly synchronized throughout the PC1 axis (Fig. 4A). Moreover, direct comparisons between *in vivo* and *in vitro* ETP, DN2a, and DN2b cells showed a strong correlation (Fig. S4A), demonstrating that pro-T cells derived from expanded HSPCs closely recapitulated the chromatin accessibility profiles of *in vivo* thymic progenitors (Supplementary Table 1).

**Figure 4.**
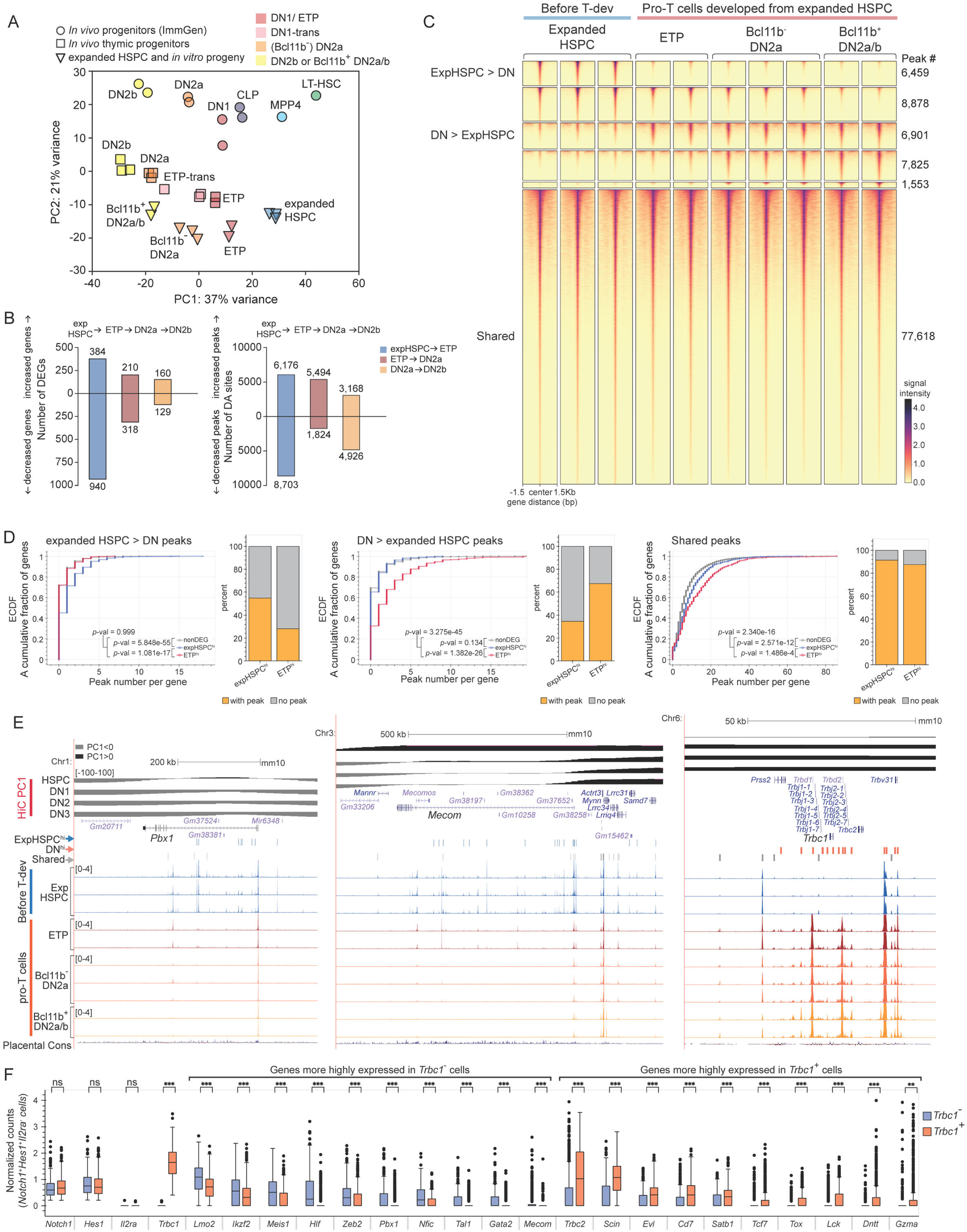
Selected developmental chromatin accessibility changes coordinated with initiating steps in T-program entry. **(A)** ATAC profiles of expanded HSPC (triangle symbols) were compared with ATAC features of *in vivo* bone marrow progenitors, *in vivo* thymic progenitors from previously published data (circle symbols)(*22*), and newly generated *in vivo* thymic progenitors’ ATAC-seq (square symbols). The principal component analysis illustrates variances of chromatin accessibility features across different progenitors. **(B)** Changes in developmental gene expression (left) and chromatin accessibility (right) across the three indicated transitions are shown. Bar graphs show the numbers of DEGs and differential accessibility (DA) sites during each developmental transition. **(C)** The peak-centered heatmap illustrates the intensity of ATAC-signal by color map in 2-3 independent replicates of expanded HSPCs and their progeny pro-T cells. The numbers of peaks in developmentally dynamic regions and stably open regions are noted. **(D)** The number of ATAC peaks in three cell type preferential classes, tested for association with genes showing the indicated patterns of developmental gene expression (key) in expanded HSPCs compared with their descendent *in vitro* ETPs. Empirical cumulative distribution functions (ECDF) enumerate the fractions of genes showing the indicated numbers of peaks per gene. Note that values for non-DEG associated peaks (gray) fall directly underneath the red curve in the left panel and under the blue curve in the middle panel. Bar graphs depict the percentage of genes harboring at least one peak vs. no peak. Adjusted *p-*values were calculated to compare distributions using a two-sample Kolmogorov-Smirnov test. **(E)** Representative UCSC genome browser tracks display stage-preferential open chromatin regions and shared open chromatin regions near *Pbx1*, *Mecom*, and *Trbc* genes. 3D chromatin states determined by previously reported HiC results (*23*) were displayed using PC1 values, in which positive values indicate active A compartment (black color near the top) and negative values correspond to inactive B compartment (gray color near the top). The expanded HSPC preferential (blue bar), DN preferential (orange bar), and shared (gray bar) peaks were noted above the tracks. **(F)** The expression levels of indicated genes were compared between *Trbc1*^-^ vs. *Trbc1*^+^ subsets of Notch-signaled ETP cells. A differential expression test was performed using the non-parametric Wilcoxon rank sum test. *** = adj *p-*value < 0.001 and absolute fold-change ≥ 1.5, ** = adj *p-*value < 0.01 and absolute fold-change ≥ 1.5, ns = not significant.

We clustered discrete genomic regions according to their dynamic chromatin accessibility patterns as expanded HSPCs progressed through early T-developmental steps (Fig. 4B), identifying specific sites that gained or lost accessibility. However, among 6 distinct ATAC site groups, the largest site group by far (∼75%) consisted of the open chromatin regions that were unchanged across the expanded HSPC stage to the DN2b stages (Fig. 4C, “Shared” group, ∼77,600 peaks). This group of sites not only included regulatory regions of common hematopoietic lineage genes (e.g., *Runx1* and *Runx3*) but also key genes establishing T cell programs, including *Tcf7*, *Gata3*, *Notch1,* and *Ikzf1* (Supplementary Table 3). Strikingly, some of the genes expressed only during T cell development, such as *Cd3g/Cd3d/Cd3e* cluster genes and distal *Trbv* regions, themselves harbored “shared” ATAC sites that were already fully accessible within expanded HSPCs and *in vivo* bone marrow progenitor populations before starting the T cell program, and these sites remained open in all of their DN descendants (Fig. S4B, C, Supplementary Table 3).

Of the developmentally dynamic chromatin accessibility patterns, the earliest changes associated with initiation of the T-cell program were of particular interest, i.e., the ATAC changes from expanded HSPCs to the early DN cells (ETP and DN2a Phase 1 pro-T cells; Fig. 4C). Notably, ∼6,500 sites substantially reduced their chromatin accessibility (Fig. 4C, expanded HSPC > DN group), whereas another ∼7,000 sites increased chromatin accessibility after entering the pro-T cell stages (Fig. 4C, expanded DN > HSPC group). These dynamic chromatin regions were highly associated with transcriptional changes. The genes more highly expressed in expanded HSPCs (expHSPC^hi^ DEGs) before Notch signaling were significantly enriched for linkage to HSPC > DN ATAC peaks, whereas the genes that were induced in ETPs (ETP^hi^ DEGs), were instead enriched for linkage to DN > HSPC ATAC peaks (Fig. 4D; see legend for non-DEGs). For instance, sharply reduced ATAC signals were often observed near LT-HSC signature genes (e.g, *Tcf15* and *Mecom*), which are turned off as expanded HSPCs transition to the ETP DN stage (Fig. 4E), while DN-preferential chromatin opening sites appeared near genes that are upregulated in pro-T cells starting from the ETP stage (e.g., *Gata3*, *Tcf7*, *Satb1*, *Dtx1*, *Gzma*, and *Lck,* Fig. S4C, Supplementary Table 3).

The expanded HSPC > DN group sites were enriched for ETS and Runx motifs, along with enriched motifs for GATA factors, bZIP family members (e.g. Jun), and C/EBP factors (Fig. S4D). While regions in the expanded HSPC < DN group were also enriched for ETS and Runx factor motifs, the sites newly opening in the DN pro-T cells were additionally enriched for bHLH factor, IRF factor, and Meis1 target motifs (Fig. S4D). These shared and differential motif patterns within dynamic chromatin regions thus indicate possible points of interaction between shared factors, like ETS and Runx factors, and context-specific factors for coordinating the earliest developmental events.

### Initiating steps in T-lineage entry marked by TCRβ gene complex activation

Of note, sites in the germ-line TCRβ regions (*Trbj-Trbc* cluster) showed a remarkable chromatin accessibility increase starting from the ETP stage (Fig. 4E). Transcription from the unrearranged TCRβ locus (*Trbc1, Trbc2*), presumably from a DJβ-associated promoter (*41*), was found in all the ETP clusters in scRNA-seq (Fig. 3D, Fig. S4E), as has been noted in ETPs before. However, the sharp increase in chromatin accessibility here suggested that its onset of expression could be a discrete event marking an initial step of the T-cell program, perhaps in response to strong Notch signaling. Notch signals are known to be required for T-lineage initiation; indeed, over 85% of *Trbc1*-expressing ETP cells (*Il2ra*^-^ cells at d 5 in our scRNA-seq analysis) showed evidence of effective Notch response (Notch1^+^ and/or Hes1^+^) (Fig. S4F). However, Notch signals are not sufficient for T-lineage initiation as they also govern numerous other developmental choices. To gain insight into any regulatory differences besides Notch signaling that could influence T-lineage entry kinetics, we closely examined the *Notch1*^+^*Hes1*^+^ *Il2ra*^-^ cells (Notch-signaled ETPs) in day 5 DNs to assess any additional features that distinguished the cells that had turned on *Trbc1* from those that had not.

Indeed, a minority of this Notch-signaled population had not turned on *Trbc1* (Fig, S4G, 42.85% of fresh progenitor progeny and 13.58% of expanded HSPC progeny), although these *Trbc1^-^* cells were in the same clusters interspersed with *Trbc1^+^* cells in UMAP visualization (Fig. S4G). Pseudobulk comparisons of these subsets of Notch-signaled ETPs revealed that the genes more highly expressed in *Trbc1*^-^ cells than in *Trbc1^+^* cells included documented regulators of progenitor genes and non-T cell genes, specifically *Lmo2*, *Meis1*, *Pbx1, Gata2*, *Mecom, Dach1* (*42*), and *Tfec*, and stem-cell markers *Fgd5* (*43*) and *Pdzk1ip1* (*44*). These genes were also closely associated with the small group of ATAC sites preferentially seen in expanded HSPC and lost in DN cells (Fig. S4H). In contrast, the genes with increased expression in the *Trbc1*^+^ ETP cells included *Tcf7*, *Lck*, *Tox*, *Satb1*, and *Dntt,* all associated with the T-lineage program (Fig. 4F, Supplementary Table 4). These results suggest that the earliest event distinguishing entry to the T cell pathway involves downregulating select transcription factors that contribute to the stem and progenitor programs and alternative lineage potentials while beginning to activate *Tcf7, Satb1, Tox,* and a select group of other lymphoid genes, even before larger transcriptome differences are manifest.

### Flt3L priming assists expanded HSPCs to initiate T-development without altering baseline gene expression or chromatin accessibility profiles

To explore how environmental signaling conditions impact this system, we tested whether additional cytokines/chemokines could influence the initial speed of developmental progression of expanded HSPCs towards the T cell fate. The thymic seeding populations (TSPs) include lymphoid-primed multipotent precursors (LMPPs) and common lymphoid progenitors (CLPs) (*5, 7, 8, 45*). Both are characterized by expression of cytokine and chemokine receptors that are genetically essential for thymic seeding *in vivo* (*46–49*). Consequently, we hypothesized that select chemokines/cytokines might provide lymphoid-lineage priming effects on expanded HSPCs. As lymphoid-lineage primed bone marrow progenitor cells express Flt3 and/or IL-7 receptor, we tested Flt3-ligand (Flt3L) and IL-7 (*8, 45, 49*), along with chemokine CXCL12, which could have an instructive role as it is required for thymic homing in HSPCs (*50*). To validate these possibilities, ELSK-sorted expanded HSPCs were treated with Flt3L, IL-7, or CXCL12 singly or in combination during the last 4-5 days of the expansion period and then transferred to an *in vitro* T-development culture (Fig. 5A).

**Figure 5.**
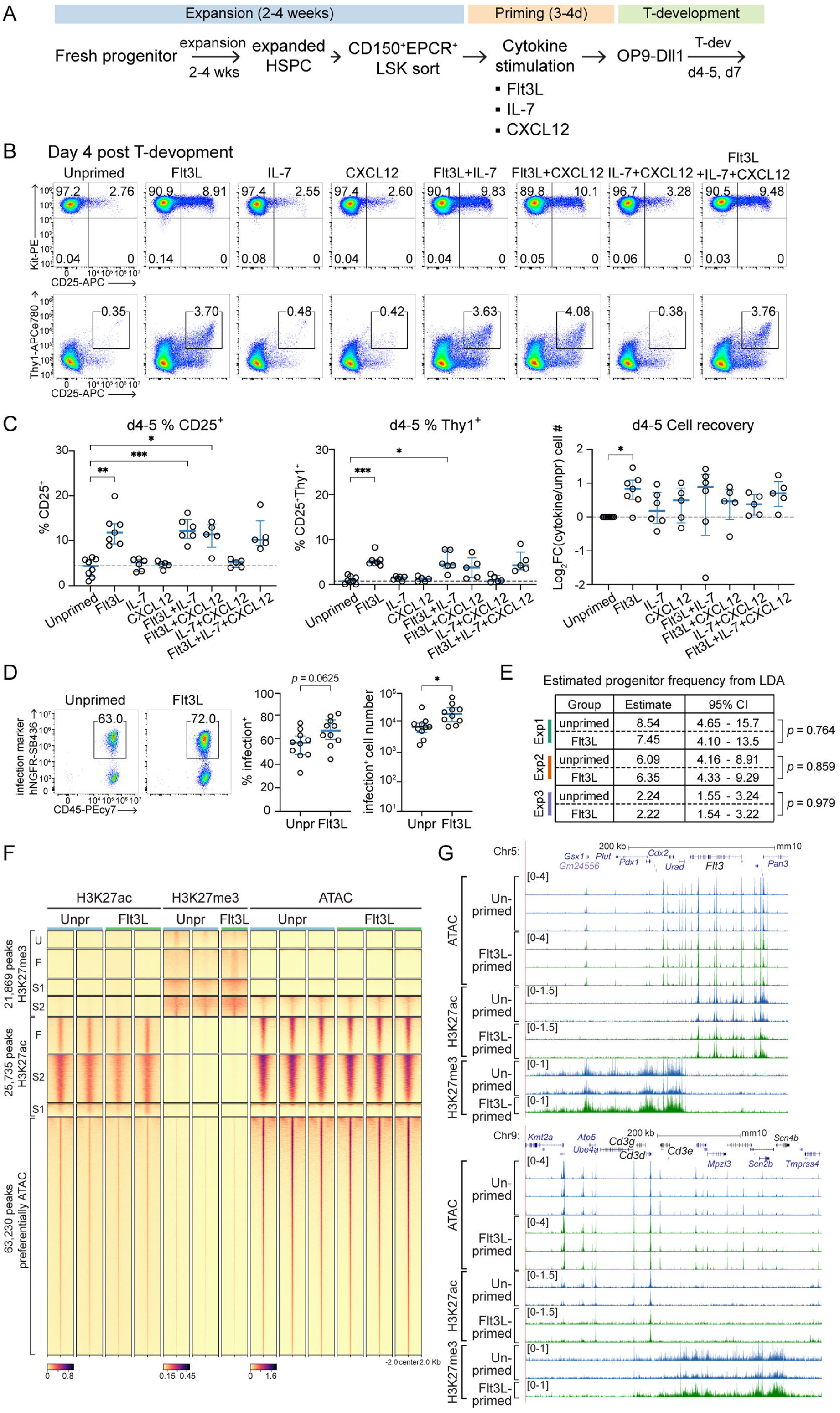
Flt3L pre-treatment enhances entry to the T cell program. **(A)** Experimental design to test the effects of different cytokine and chemokine stimulation on early T cell development. **(B, C)** Expanded HSPCs were treated with Flt3L, IL-7, or CXCL12 singly or in combination prior to OP9-Dll1 co-culture. T-development progression was assessed using flow cytometry for Kit, CD25, and Thy1 expression, and cell recovery was compared. Representative flow cytometry plots (B) and the results from 5-6 independent experiments are shown with median values and interquartile ranges (C). The dotted line indicates the median value of unprimed conditions. Statistical comparisons by one-way ANOVA. *** = *p-*value < 0.001, ** = *p-* value < 0.01, * = *p-*value < 0.05. **(D)** Expanded HSPCs were treated with Flt3L for 4 days, infected with retrovirus expressing human NGFR, and subjected to T cell development. Infection rates and infected cell numbers of unprimed vs. Flt3L-primed progeny were displayed. Unpaired t-test. * = *p-*value < 0.05. **(E)** The frequencies of T cell progenitors in expanded HSPCs that were either unprimed or Flt3L-primed were estimated using three independent limiting dilution assays. **(F)** The peak-centered heatmap illustrates the patterns of H3K27ac and H3K27me3 (measured by CUT&RUN) and chromatin accessibility profiles (measured by ATAC) in unprimed and Flt3L-primed HSPCs. U: unprimed > Flt3L-primed, F: Flt3L-primed > unprimed, S1: shared and closed sites, S2: shared and open sites. **(G)** Representative UCSC Genome Browser tracks depict the features of ATAC, H3K27ac, and H3K27me3 in unprimed (blue) and Flt3L-primed (green) HSPCs near the *Flt3* and *Cd3* genes.

Notably, among 8 different cytokine and chemokine conditions, pre-treatment with Flt3L achieved faster ETP to DN2 transition (determined by CD25 upregulation) starting from day 4 (Fig.5B, C), before *Bcl11b* was upregulated (Fig. S5A). Flt3L alone produced the maximum effect, with no further additive effects by combination with IL-7 or CXCL12 (Fig. 5B, C), which were also ineffective alone. The Flt3L pre-treated cells also accelerated their upregulation of Thy1 and Bcl11b by day 7, demonstrating that such Flt3L pre-stimulation primed expanded HSPCs for faster T-cell differentiation through DN2a to DN2b stages. (Fig. S5B, C). Moreover, Flt3L-primed cells resulted in superior cell recovery compared to unprimed controls after retroviral transduction (Fig. 5C, D, S5C).

Flt3L signaling via Flt3 could alter the readiness of individual cells to enter the T-cell pathway; alternatively, it could preferentially stimulate outgrowth of a pre-existing subpopulation of expanded HSPC cells with enhanced T-lineage potential. In the latter case, the Flt3L-primed cells should exhibit a higher pro-T-cell precursor frequency.

However in repeated limiting dilution analyses, both unprimed and Flt3L-primed expanded HSPCs had comparable T-precursor frequencies in independent trials giving values ranging from averages of 1/2.2 (unprimed) vs. 1/2.2 (Flt3L-primed) to averages of 1/8.5 (unprimed) vs. 1/7.5 (Flt3L-primed) (Fig. 5E). This suggests that Flt3L pre-treatment does not select for a subset of cells with increased T cell potential, but rather alters cells across the population as a whole to increase their pro-T cell developmental speed.

To investigate whether Flt3L pre-treatment alters the pathway through which expanded HSPCs reach the DN2a and DN2b stages, we compared transcriptional profiles, chromatin accessibility patterns, and active (H3K27ac) and repressive (H3K27me3) histone marks between Flt3L-primed and unprimed expanded HSPCs (Fig. 5F). Surprisingly, Flt3L treatment did not detectably alter gene expression patterns in the expanded precursors despite enabling the cells to progress significantly faster during pro-T cell stages once triggered by Notch signaling. In particular, it showed no evidence of changes anticipating the *Trbc1^-^* to *Trbc1^+^* regulatory shifts noted above (Fig. S5D). Similarly, when ELSK-purified expanded HSPCs were split into unprimed and Flt3L-primed subcultures for 4 days, the ATAC-seq profiles and histone marks from these subcultures showed minimal differences globally (Fig. 5F, S5D).

Developmentally regulated regions such as *Flt3* and *Cd3* gene cluster regions also showed closely matching chromatin features between Flt3L-stimulated expanded HSPCs and unstimulated expanded HSPCs (Fig. 5G). Although Flt3L-stimulation in expanded HSPCs did not show obvious gene expression or chromatin feature changes at the bulk population level, scRNA-seq results detected a modest but broad stress response to Flt3L treatment, represented by increased expression of *Fos*, *Egr1*, *Txnip*, and *Bnip3* (Fig. S5E). These results suggest that Flt3L/Flt3 signaling in expanded HSPCS does not work through early activation of lymphoid gene expression or chromatin program, but probably by priming cells to be readily activated or differentiated.

In single-cell transcriptome analysis, pro-T cells derived from Flt3L-treated expanded HSPCs showed slightly faster upregulation of DN2/DN3 signature genes, such as *Bcl11b*, *Trgc-C2*, *Il2ra*, *Gzma*, and *Trdc*, and downregulation of progenitor genes *Hoxa9* and *Mef2c* (Fig. S5E). However, the progeny of Flt3L-primed expanded HSPCs exhibited overall gene expression programs that were nearly indistinguishable from corresponding progeny of unprimed expanded HSPCs, with highly overlapping developmental trajectories (Fig. S5F). Thus, 4-day Flt3L-priming primarily preserves normal gene expression programs in expanded HSPCs while usefully preparing cells to respond to T-cell inductive signaling with faster developmental kinetics.

### Dissecting roles of 23 transcription factors in early T cell development using CRISPR-Cas9 mediated acute deletion

The full characterization of the expanded HSPC system enabled us to now leverage this system to dissect the roles of progenitor- and lymphoid-biased regulators at the outset of T-cell development. Previous studies have identified the roles of some of these factors in early T cells; for example, Runx1 has been shown to act as a T-lineage accelerator (*51*), while *Spi1* and *Erg* have been shown to act as T-lineage brakes (*37, 52*). However, the precise functions of many other factors are poorly understood or remain uncharacterized outside of leukemic contexts. We reasoned that the extended duration of early pre-commitment stages in expanded HSPC-derived pro-T cells could provide a unique opportunity to elucidate the roles of less explored transcription factors during the early ETP stages.

To this end, we selected 23 transcription factors expressed in HSPCs and/or ETPs (Fig. S6A) and assessed their roles by acutely disrupting them with the CRISPR-Cas9 system (Fig. 6A). These factors fell into four groups: (i) genes encoding the stem and progenitor cell regulators that bridge the prethymic-to-early intrathymic transition, some of which have known roles in leukemia (*Hoxa9*, *Meis1*, *Lmo2*, *Tal1*, *Lyl1*, *Spi1*, *Hlf*, *Hmga2*); (ii) genes encoding known T-lineage essential targets (*Ikzf1*, *Myb*, *Hes1*, *Tox*); (iii) genes encoding the immediate-early activation factors Fos and FosB; (iv) genes encoding possible interacting partners of the widely-used factor Runx1, which could elucidate how Runx1 accelerates T-lineage program entry specifically (*51*). Considering that Runx occupancy sites are commonly enriched for ETS motifs along with Runx motifs (*39*), we surveyed multiple genes encoding ETS family transcription factors that are implicated as potential partners for Runx1 (*Ets1*, *Ets2*, *Fli1*, *Gabpa*, *Erg*, *Etv6*, *Elf1*, *Elk3*, *Elk4*) as well as PU.1 (*Spi1*), a known Runx1 partner (*51, 53*). For each target transcription factor, we designed 2-3 distinct guide RNAs (gRNAs), which were introduced by retroviral transduction into expanded HSPCs, either primed with Flt3L or untreated (Fig. 6A). The gRNA-infected HSPCs were then co-cultured with OP9-Dll1 cells to evaluate their progression through early T cell development by measuring CD25 and Thy1 induction in cells positive for the RV infection marker. The impact on expression of these stage-progression marker genes is shown in Fig. 6B; Fig. S6A. Additionally, we compared cell recoveries to determine how the loss of target transcription factors impacted cell survival and proliferation program (Fig. 6C). Of note, we utilized HSPCs expressing a *Bcl2* transgene to minimize survival defects possibly caused by a loss of a factor; nevertheless, some strong cell recovery impacts were seen. A summary of the results for differentiation and cell recovery, respectively, for each perturbation is given as a set of heat maps in Fig. 6D, and examples of the phenotypes observed are shown in Fig. 6E and Fig. S6B.

**Figure 6.**
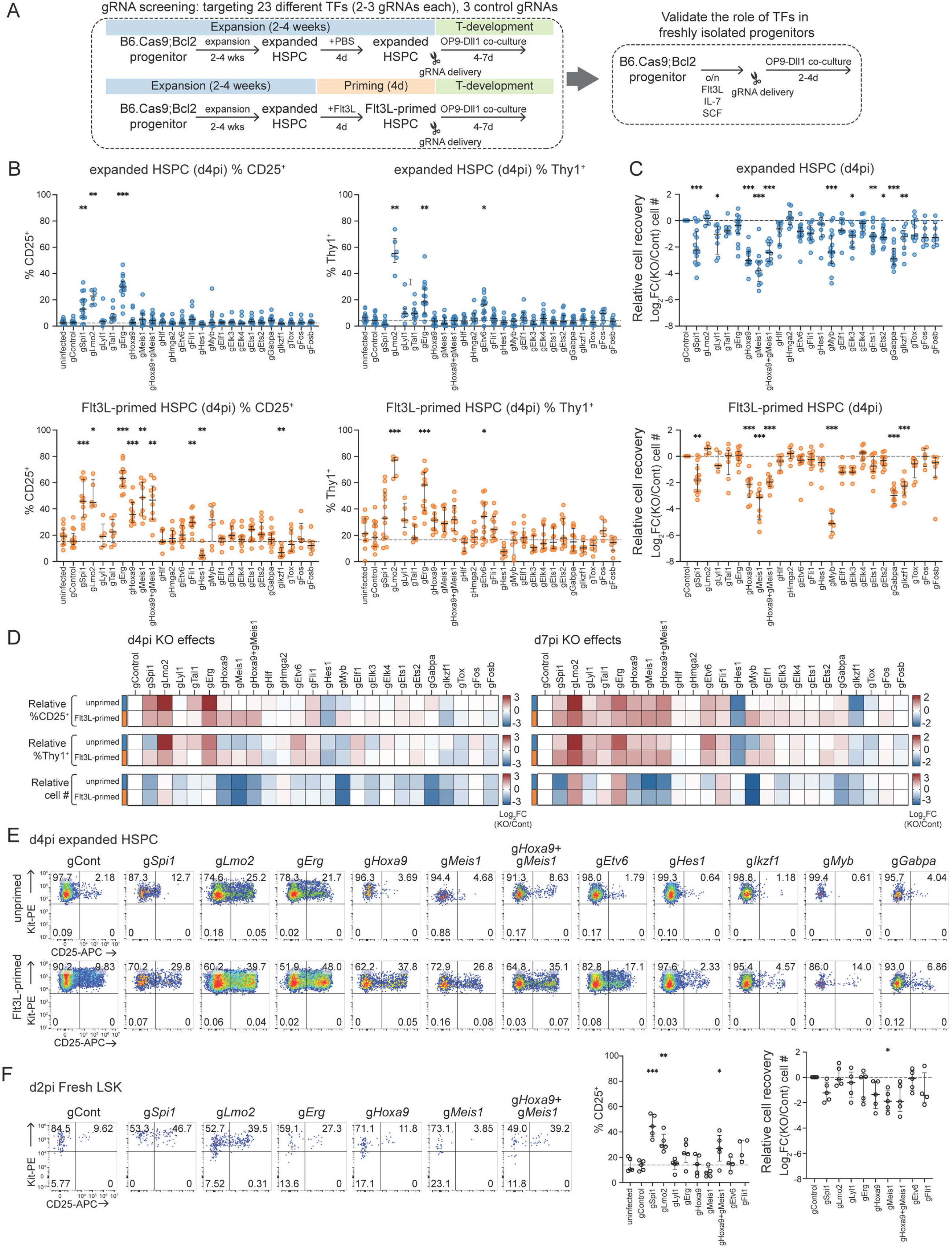
Acute disruption of early T cell transcription factors reveals the roles of PU.1, Lmo2, Erg, Hoxa9, and Meis1 in developmental progression. **(A)** Experimental schematic describes CRISPR-Cas9 screening and validation strategy to identify transcription factors (TFs) controlling early T cell development. **(B-E)** gRNAs against 23 different TFs were introduced to the expanded HSPCs without or with Flt3L priming, and the *in vitro* early T cell development was induced using OP9-Dll1 co-culture. After 4 or 7 days post-OP9-Dll1 co-culture, T-developmental progression was assessed by measuring CD25, Thy1, and Kit expression levels (B, D). Relative cell recovery was evaluated by comparing cell number to that of the control conditions (C, D). **(B, C)** The graphs summarize the results from 5-14 independent experiments with median values and interquartile ranges. Individual data points represent the mean of 1-3 technical replicates. The dotted line indicates the median values of the control group. Brown-Forsythe and Welch ANOVA test with Dunnett’s T3 multiple comparisons (B) and Kruskal–Wallis test (C) for the statistical test. * = *p-*value < 0.05, ** = *p-*value < 0.01, *** = *p-*value < 0.001. **(D)** Heatmaps illustrate the fold-change (FC) of the frequencies of CD25^+^, and Thy1^+^ cells, and cell numbers relative to the control conditions. The color intensity depicts the indicated changes in Log_2_(FC+0.01). **(E)** Representative flow cytometry plots demonstrate the effect of loss of indicated TFs in T cell development determined by Kit and CD25 levels. The perturbation effects in unprimed expanded HSPC and Flt3L-primed expanded HSPCs were measured at 4 days post-infection. **(F)** Graphs present the validation test of loss of indicated TFs in fresh LSK progenitors at day 2 post-infection. The frequency of CD25^+^ cells and relative cell number by comparing with control conditions were shown with median and interquartile ranges. The dotted line indicates the median values of the control group. Ordinary one-way ANOVA with Dunnett’s T3 multiple comparisons. * = *p-*value < 0.05, ** = *p-*value < 0.01, *** = *p-*value < 0.001.

To explore the observed phenotypes, we first focused on the deletion of the PU.1 coding gene, *Spi1*, as its role in the DN1 stage has been extensively studied (*37, 54*). Acute disruption of *Spi1* caused significant cell losses despite the presence of the *Bcl2* transgene, as previously reported (Fig. 6C), but significantly accelerated the induction of CD25 and Thy1 among the surviving cells from both unprimed expanded HSPCs and Flt3L-primed expanded HSPCs (Fig. 6B, Fig. S6C, D). These results confirmed that PU.1’s established role in slowing T-cell developmental progression was faithfully captured by the expanded HSPC system.

Among the genes with known positive roles in T cell development, *Hes1*, *Ikzf1*, *Myb*, and ETS family transcription factor gene *Gabpa* caused notable phenotypes. Loss of *Myb*, *Ikzf1,* or *Gabpa* resulted in catastrophic drops in cell recovery despite the *Bcl2* transgenic background of the cells, with differentiation of the survivors only detectably inhibited by loss of *Ikzf1*. In contrast, loss of *Hes1* had little effect on cell yield, but selectively inhibited developmental progression (a fourfold drop in % of CD25^+^ cells).

Although most of the knockout factors had little significant functional impact on T cell developmental progression, a select set of bone marrow progenitor-inherited factors were inferred to have substantial regulatory roles. This included Lmo2, Hoxa9, and Meis1, as well as the ETS family member, Erg. A weaker effect was also seen for Etv6. All these TFs are normally repressed after T-lineage commitment (Fig. S6A), otherwise their normal endogenous levels hinder some aspects of early T-developmental progression. However, each TF appears to delay T-developmental progression by targeting distinct sets of programs, as suggested by previous transcriptome analyses of a smaller set of factors (*55*). Here, some differences could be seen as they reduced the normal concordance between CD25 vs. Thy1 upregulation (Fig. S6C). Further exploration of the prolonged ETP stage in expanded HSPCs could give much needed experimental access to the distinct networks controlled by these factors, as shown for Lmo2 below.

Overall, the effects on differentiation speed and the effects on population growth appeared to be differentially regulated across the knockout factors (Fig. 6D). Accelerated differentiation in the cases of *Spi1* and *Hoxa9* and/or *Meis1* knockouts was complicated by the severe population losses that these perturbations inflicted, comparable to the cell losses seen when T-lineage essential regulatory genes *Ikzf1*, *Myb*, and *Gabpa* were deleted. In contrast, losses of *Erg* and *Lmo2* accelerated T-lineage differentiation speed without any detriment to population growth.

The abundant cells and relatively leisurely development of the expanded HSPC-derived cultures enabled numerous effects to stand out robustly (Figs. 6B, C), especially when using Flt3L-primed input cells. We confirmed the strongest of these effects in freshly isolated LSKs as well, by examining early (day 2 and day 4) timepoints after gRNA transduction (Fig. 6F, S6E): loss of *Spi1* or *Lmo2* alone accelerated progression to DN2 stages in day 2 fresh LSK cultures, and although *Meis1* or *Hoxa9* single knockouts did not cross the significance threshold, the combined knockout was effective (Fig. 6F). Thus, the expanded HSPC system is a sensitized, kinetically favorable experimental framework in which to dissect the roles of transcriptional regulators in the transition from prethymic to early intrathymic T-cell precursors.

### Acute disruption of *Lmo2* in expanded HSPCs reveals its previously uncaptured role in regulating T cell developmental kinetics

Lmo2 and Erg stood out as factors that were dispensable for viability and cell recovery but clearly exerted restraint of progression to the DN2 stage at their normal levels of expression. In previous work, we reported that although disruption of *Erg* allowed faster upregulation of some DN2-associated genes, it actually caused a deviation from the normal T-lineage gene expression program (*37*), suggesting that Erg normally has a lineage fidelity-promoting role despite its slowing of T-cell development and being dispensable for population size. This could be consistent with *Erg*’s continued expression into DN3a stage (*56, 57*). *Lmo2*, on the other hand, is turned off in the early ETP stage soon after cells enter the thymus, and its abnormal expression at later stages is invariably associated with T-ALL (*58, 59*). To explore how endogenous Lmo2 normally interacts to control the speed of the T cell differentiation mechanism, we determined how its loss affected the T-differentiation gene expression program.

First, to resolve the initiation and persistence of the acceleration phenotype observed in the *Lmo2* knockout (KO) (Fig 6E), we analyzed multiple timepoints for developmental progression by cell surface markers CD44 and CD25 in the progeny of expanded HSPCs infected with control vs. *Lmo2* KO vectors, either with or without Flt3L-priming (Fig 7A). Here, CD44^+^ CD25^-^ phenotype defined DN1 cells, which include ETPs. *Lmo2* KO noticeably influenced developmental progression as early as day 3, where there was 4-5× higher frequency of the DN2 population observed in the progeny of expanded HSPCs with the *Lmo2* KO vs control (Fig 7B), indicating that Lmo2’s influence on T cell developmental kinetics occurs at the earliest timepoints. Thy1 was upregulated even earlier than CD25 (Fig S7B), showing that the DN1 cells were affected beyond the *Il2ra* gene encoding CD25 itself. Furthermore, the acceleration phenotype continued past the DN2 stage: on day 8 of co-culture with unprimed expanded HSPCs, ∼70% of progeny with the *Lmo2* KO reached the DN3 stage while ∼85% of progeny of control-infections remained in the DN1 stage (Fig 7B). Thus, the accelerated DN1-DN2b transition also maintained the appropriate conditions for continuing development through the DN2-DN3 transition. Acute disruption of *Lmo2* in expanded HSPCs increased the proportion of progeny occupying later DN stages at every subsequent timepoint (Fig. 7C), regardless of priming conditions. Though Flt3L-primed infected samples yielded even higher frequencies of progeny at later DN stages at every timepoint, the Flt3L-priming effect was greatly overshadowed by the impact of the *Lmo2* KO itself (Fig S7A). Thus, the mechanisms that launch the T cell program due to loss of Lmo2 are separable from those involved with the Flt3L-priming effect.

**Figure 7.**
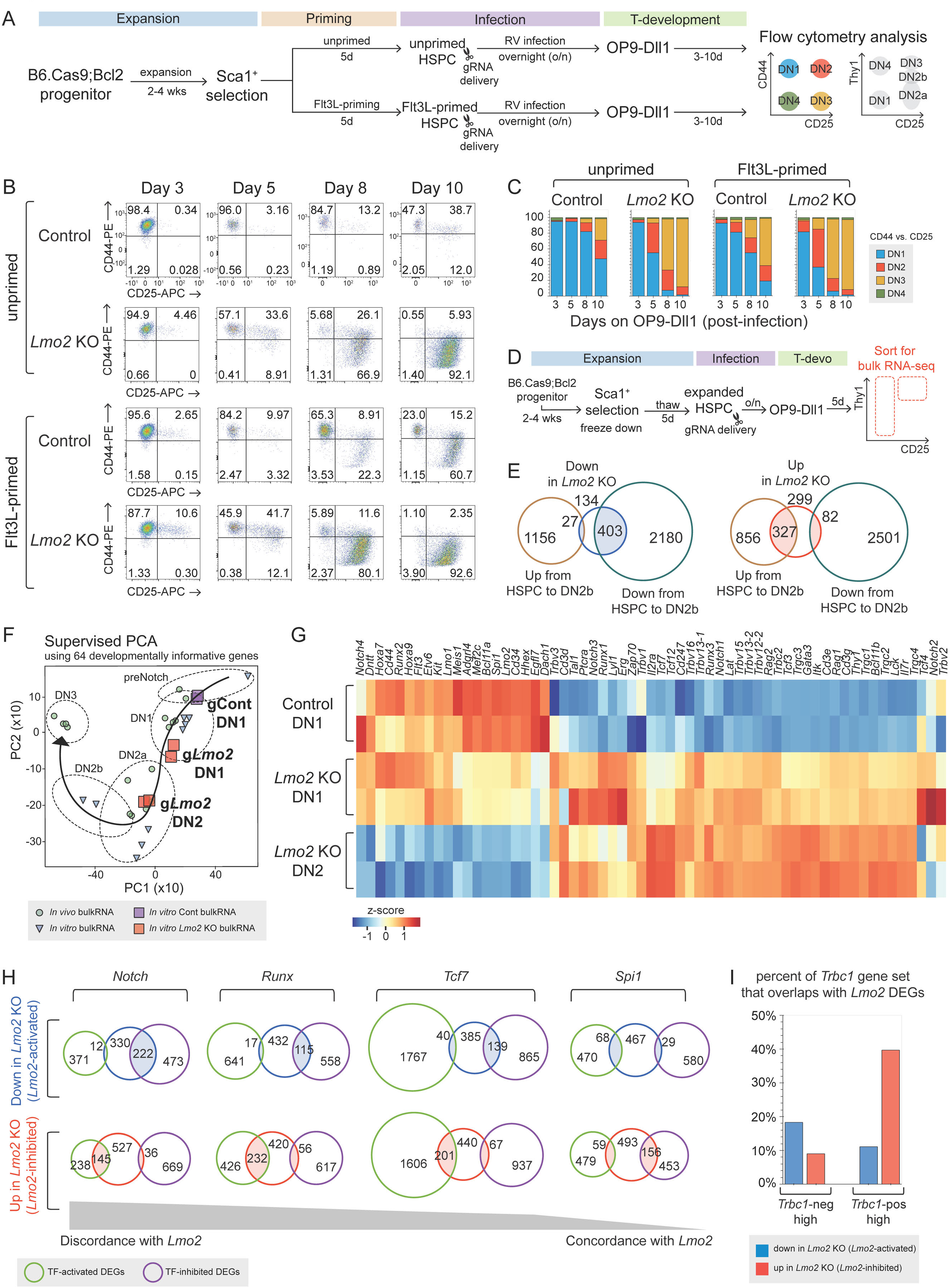
Loss of Lmo2 results in accelerated induction of the T cell program, partially consistent with Notch and Runx transcriptional activity. **(A)** Experimental design for analyzing *in vitro* OP9-Dll1 co-cultures seeded with control vs. *Lmo2* KO expanded HSPCs, either unprimed or Flt3L-primed, across multiple timepoints. **(B, C)** Representative flow plots showing DN1-4 frequencies by CD44 and CD25 for each timepoint and condition (B). Stacked bar plots quantitatively visualize the average frequencies from 2 independent experiments with 3-4 technical replicates (C). **(D)** Experimental design for comparing transcriptional profiles of control vs *Lmo2* KO progeny of unprimed expanded HSPCs with bulk RNA-seq. **(E)** Venn diagrams compare DEGs between Cont-DN1 and *Lmo2*KO-DN1 populations from 2 independent experiments. DEGs between HSPC and DN2b cells are generated as described in Fig 2. DEGs were defined using adj *p*-value < 0.05 and absolute log2FC > 0.5. **(F)** Supervised PCA plot using 64 T-developmental trajectory genes shows control and *Lmo2* KO pro-T cells mapped along with reference data, as described in Fig 2E. **(G)** Heatmap illustrates the *Z*-score of normalized TPM values for select genes across all replicates. **(H)** Venn diagrams compare *Lmo2* DEGs ( “*Lmo2*-inhibited”: upregulated in *Lmo2* KO, “*Lmo2*-activated”: downregulated in *Lmo2* KO) to published DEG lists for *Notch, Tcf7, Runx* and *Spi1*. Order of comparisons is defined by discordant vs. concordant activity. **(I)** Bar plots show the percent of DEGs enriched in *Trbc1*^-^ ETP or *Trbc1*^+^ ETPs that overlap with *Lmo2*-inhibited or *Lmo2*-activated target genes.

Of note, *Lmo2* KO did not cause survival defects in unprimed or Flt3L-primed samples, confirming results seen in the CRISPR/Cas9 screen (FigS7C); thus, observed kinetic differences were not caused by a selection artifact. Importantly, the acceleration phenotype driven by acute knockout of *Lmo2* was also repeatedly observed in faster-differentiating Lin^-^ progenitors from fresh bone marrow, provided that *Lmo2* was efficiently disrupted by the outset of the OP9-Dll1 culture. However, these fresh progenitors allowed a smaller window of time for comparison and a smaller fold change: lin^-^ bone marrow progenitors with *Lmo2* KO had ∼65% of their progeny in the DN2 stage by day 3, as compared with 35% in controls (Fig S7D). Taken together, these results supported the significance of Lmo2 as a kinetic block on T cell development, acting at the earliest timepoints right as progenitors encounter Notch signaling, and the expanded HSPCs presented a novel opportunity to fully explore the earliest-acting effects of Lmo2, previously inaccessible for study.

We next asked how the loss of Lmo2 causes developmental acceleration. Two non-exclusive explanations were considered: (1) loss of Lmo2 prematurely starts the normal sequence of developmental events, only affecting the rate at which these events are initiated and thus suggesting that downregulation of Lmo2 is a bottleneck for the upregulation of the T cell program; or (2) loss of Lmo2 allows access to an alternate trajectory that bypasses the normal steps in T cell development but ultimately arrives at the same destination, thus suggesting that Lmo2 normally blocks this alternate route to maintain fidelity to the canonical pathway. To resolve whether *Lmo2* KO causes developmental acceleration along canonical T cell developmental pathways or alternate pathways, we thus investigated the immediate transcriptional events caused by loss of Lmo2 with bulk RNA-seq for sorted populations on day 5 of OP9-Dll1 co-culture, the earliest timepoint where a large acceleration phenotype is observed in *Lmo2* KO (Fig 7D, Fig S7A). The sorted populations at this timepoint include DN1 cells from control-infections (Cont-DN1) and DN1/DN2 cells from infections with *Lmo2* KO vectors, sorted by Thy1, CD25 and CD44 (*Lmo2*KO-DN1, *Lmo2*KO-DN2).

Despite their similar cell surface phenotypes, Cont-DN1 cells and *Lmo2*KO-DN1 cells already showed major transcriptional differences at this timepoint, with DEG analysis revealing 564 DEGs downregulated due to Lmo2 knockout, and 708 DEGs upregulated due to *Lmo2* KO (padj < 0.05, absolute log2FC > 0.5). To assess whether these transcriptional differences are relevant to normal T cell development, we compared the *Lmo2* KO DEGs to DEGs generated between reference uninfected expanded HSPCs and uninfected expanded HSPCs co-cultured on OP9-Dll1 to the DN2b stage. Fig 7E shows that the *Lmo2*-activated DEGs, downregulated by the *Lmo2* KO, greatly overlapped with the DEGs that are normally downregulated from HSPC to DN2b; similarly, the *Lmo2*-inhibited DEGs, upregulated by the *Lmo2* KO, greatly overlapped with the DEGs normally upregulated from HSPC to DN2b. Thus, the genes either directly or indirectly influenced by Lmo2 highly corresponded with the genes that are normally upregulated/downregulated throughout normal T cell development. This is a significant contrast with the effect of the *Erg* knockout, which accelerated development but biased the cells towards an alternate trajectory, reflected by the poor overlap between *Erg*-related DEGs and the HSPC-to-DN2b DEGs (Fig S7E). The *Lmo2* KO seemed to accelerate the T-cell program specifically: svPCA analysis mapped *Lmo2*KO-DN1s (large red boxes) closer to DN2a references, while Cont-DN1s (large purple boxes) mapped closer to early DN1 references (Fig 7F). Interestingly, *Lmo2*KO-DN2s (large red boxes), which were already abundant by day 5 of co-culture, mapped neatly within the normal DN2a/DN2b reference points, so their DN2 cell surface phenotype now faithfully reflected their precocious differentiation on a transcriptional level.

### *Lmo2* deletion unleashes a broad spectrum of T-cell gene expression

These analyses that integrated reference populations supported the hypothesis that Lmo2 normally slows entry to the canonical T cell pathway, rather than blocking an alternate trajectory towards the T cell fate. Interestingly, though, comparison of normalized TPM values just between Cont-DN1, *Lmo2*KO-DN1 and *Lmo2*KO-DN2 showed an important abnormality: the T cell program genes were upregulated even before downregulation of key stem/progenitor program members (Fig 7G). *Lmo2*KO-DN1 cells retained high expression of stem/progenitor members, with *Hoxa9, Hoxa7* and *Flt3* at comparable levels of expression to Cont-DN1s, and *Meis1, Mef2c, Bcl11a, Spi1* and *Hhex* at only slightly lower levels of expression than Cont-DN1s but still much higher than *Lmo2*KO-DN2. Concurrently, *Lmo2*KO-DN1 cells highly expressed T cell program TFs *Tcf7, Gata3,* and *Bcl11b* (normally only activated in DN2 cells) and TCR signaling genes including *Lat, Itk, Lck, Zap70, Cd3e*, and *Cd3g* (Fig S7G), only slightly less than the expression seen in *Lmo2*KO-DN2 cells. T lineage promoting regulators *Runx1* and *Runx3* were also included in the upregulated group, along with *Notch1, Notch2* and *Notch3*. Of note, transcription of the TCRβ-coding locus was also sharply increased in gLmo2-DN1s (*Trbv12-2, Trbv13-2, Trbv15, Trbc2* transcripts; Fig S7F)(cf. Fig. 4F, Fig. S4E-G). Thus, a robust T cell signature was activated prematurely without a full shutdown of the rest of the stem/progenitor signature, suggesting that Lmo2 itself could be the key limiting factor for T-lineage initiation.

Deletion of Lmo2 had an asymmetric effect on the positive and negative regulatory mechanisms that distinguish *Trbc1*-neg cells from *Trbc1*-pos cells under similar Notch conditions (Fig. 4). *Lmo2* deletion activated a large proportion of the genes upregulated in Trbc1^+^ cells (∼40%); however, loss of Lmo2 only inhibited ∼20% of the genes that were higher in Trbc1^-^ cells (Fig 7I). Thus, although endogenous Lmo2 is likely to restrain the pathways that activate *Trbc1* expression, it appears less important to sustain the full set of stem and progenitor factors that are normally downregulated in *Trbc1*^-^ to *Trbc1*^+^ progression.

### Lmo2 and the T cell specification gene network

Given that so many key T cell TFs were rapidly upregulated by loss of Lmo2, we considered the possibility that the observed acceleration was a downstream effect of the specific upregulation of a key T cell TF; in other words, Lmo2 might only be acting on a specific subset of the T cell program rather than on multiple elements of the program, as the bulk RNA results suggest. If so, *Lmo2* DEGs should highly correspond with T-cell TF-target genes. Therefore, three such TFs were considered: Notch, since Notch signaling strength is vital for T-cell development (*60, 61*); Runx family, since slight Runx dosage increase is known to cause acceleration (*51*); and TCF1 (encoded by *Tcf7*) which also has a role in accelerating development (*62*). All of these also upregulate *Bcl11b* (*38, 51*), the canonical T cell commitment marker, and could account for its precocious expression. Venn diagram comparisons in Fig 7H highlight the resulting comparisons, which showed that all three factors were highly biased to oppose Lmo2’s effect when targeting the same genes. For instance, Notch had the strongest discordant relationship with Lmo2, as the *Lmo2*-activated genes overlapped with 222 *Notch*-inhibited genes but only 12 *Notch*-activated genes, and conversely, *Lmo2*-inhibited genes overlapped with 145 *Notch*-activated genes but only 36 *Notch*-inhibited genes. Runx and TCF1 showed similarly pronounced discordances with Lmo2 (Fig. 7H). We also considered the possibility that Lmo2 could be restraining T cell development by its supportive regulation of *Spi1*, which causes an acceleration phenotype when knocked out (Fig. 6). Although there was a somewhat concordant relationship, as *Lmo2*-activated genes had greater overlap with *Spi1*-activated genes than with *Spi1*-inhibited genes and vice versa for *Lmo2*-inhibited genes, these overlaps were less impressive than the discordances with the T-lineage promoting genes (Fig 7H, Fig. S7H). In contrast, the effect of another T-lineage restraining factor, Erg (Fig. 6), was completely distinct, as reflected by the small overlap between *Erg* target genes and *Lmo2* DEGs (Fig S7I).

These results imply that *Lmo2* works through mechanisms that are partially and completely independent of its fellow stem/progenitor members *Spi1* and *Erg*, respectively.

## Discussion

The eligibility of progenitor cells to enter the T cell pathway is shaped by the gene regulatory networks that operate during the transition from pre-thymic progenitor cells to the initiation of the T cell program. Here, we showed that the i*n vitro-*expanded HSPCs with high progenitor frequencies for T cells provide new opportunities to closely monitor these steps, and that use of this system revealed transcription factors regulating the entry to the T cell program and pro-T cell pool size. In particular, these studies showed the extent to which a highly enriched pool of T-lineage competent precursors prior to Notch signaling already maintains a permissive chromatin configuration for initiating the T cell program, the transcriptional activation of the *Trbc1* and *Trbc2* regions, apparently from the Dβ-associated promoter (*41*), as an exceptionally early landmark of T-lineage initiation. In this system, the molecular changes associated with this step could be resolved with new clarity. Furthermore, because the kinetics of T-lineage progression are slower in this system than with fresh progenitors, the potent opposing actions of Meis1, Hoxa9, and Lmo2 against the T cell program could be revealed here with new sensitivity using an acute deletion approach. The effects of deleting these early T-development speed controllers were previously challenging to study (*37*) because they are usually downregulated too quickly during *in vitro* T cell development, when fresh multipotent progenitors first started the T cell program. With this system, the distinct responses to withdrawal of key factors could be seen and compared readily. Thus, Lmo2 could emerge as a very potent and distinctive controller of initiation of the T cell program overall.

Our results showed that the convenience of this system could be augmented by priming the cells with Flt3L for 4-5 days before exposure to T-lineage inducing conditions, which empowered the cells to progress into T-cell development more quickly upon Notch stimulation. Recent reports also suggest that Flt3L addition throughout HSPC expansion improves yield by suppressing CD41^+^ progenitor differentiation (*63, 64*). Nevertheless, the majority of cells with a stem-like EPCR^+^ CD150^+^ LSK phenotype appeared minimally affected in transcriptome or chromatin states upon 4-5 days of Flt3L priming. Thus, global activation state or other post-transcriptional cellular features besides gene regulatory identity could contribute to lymphoid pathway access. Notably, these microenvironmental speed control mechanisms may have *in vivo* significance as well. *In vivo*, thymus seeding progenitors in adult mice progress substantially more slowly from ETP to DN2 stage than they do in OP9-Dll1 coculture (*65*), and it is possible that the timing is affected by cytokine signals received in distinct intrathymic domains (*66*).

The role of Lmo2 shown here was particularly interesting. Lmo2 is often considered part of a positively regulating “pentamer” transcription factor complex with a GATA factor, E2A, SCL (Tal1) or Lyl1, and Ldb1, implicated in stem cell and erythroid development. In T-cell development, its ectopic reactivation has been associated with T-ALL of a late stage type (*58, 59*). A recent report showed that Lmo2 in EBF1-knockout pro-B cells may play a role in keeping the cells multipotent enough to enable T-cell potential, possibly including a role in priming the *Tcf7* locus for Notch-driven T-lineage expression (*67*). This is not incompatible with our results. However, we have shown here that the expression of Lmo2 then acts as a natural “starting gate”, holding back a multi-factor T-lineage program that explosively gets under way under Notch signaling, in a fraction of the time seen in controls, if Lmo2 is removed early. In contrast to Erg and PU.1, the removal of Lmo2 permits a largely normal T cell program to begin, and in contrast to PU.1, Meis1, and Hoxa9, it does so without any population size penalty.

The range of targets defined in any perturbation system depends on the baseline used for comparison, and the expanded HSPCs we analyzed in perturbation experiments were not identical to normal progenitors *in vivo*. Nevertheless, the targets affected by *Lmo2* deletion in this system suggest a biologically significant role. It is not surprising that endogenous levels of Lmo2 appeared to support expression of progenitor-associated regulatory genes including *Dach1, Hhex, Spi1, Bcl11a, Mef2c*, and itself (*42, 58, 59*). More notable was our evidence implicating normal endogenous Lmo2 levels as concertedly restraining expression of *Runx1, Runx3, Tcf7, Notch 1, Notch3, Gata3,* and genes encoding several bHLH factors, all of which were upregulated multiple times faster in a T-lineage inducing environment when the endogenous Lmo2 was removed. This was a remarkably concerted impact across the range of factors important to launch T-cell programming, with predictable downstream results like accelerated activation of other T-cell genes including *Bcl11b* (*38*). In contrast, Lmo2 removal did not appear to enhance innate-like gene expression, distinguishing it from other accelerators, including knockout effects of PU.1 and Erg (*37*) and overexpression effects of Runx1 (*51*). Could Lmo2 be an important initiation switch *in vivo*? Lmo2 levels do not appear to differ between Flt3L-primed and unprimed expanded HSPC, and as shown in Fig. S6, its levels are similar also in fresh LT-HSC through MPP4 cells from the ImmGen consortium (*22*). However, it is strikingly lower in CLPs and in the great majority of fresh ETPs alike. It is thus tempting to suggest that the drop in Lmo2 may be one of the key events *in vivo* that enables CLPs to respond to Notch signaling faster than MPP4 cells (LMPP) to generate T cells. More work will be required to show how Lmo2 exerts its effects in these cells.

For many developmental studies, iPSC technology provides a priceless way to “stop time” while cells can be genetically manipulated. The complexity of generating T lineage precursors from iPSCs and the difficulty of isolating them in high purity at early stages have made this technology less fruitful to dissect T-lineage developmental mechanisms, however, even though it has shown some promising results (*68, 69*). We show here that the PVA culture system for expanded HSPCs similarly offers a way to stop time upstream of T lineage program initiation, and to explore in new depth the genetic circuitry involved in this momentous lineage choice.

## Materials and Methods

### Study design

This study was designed to reveal 1) the earliest changes in gene expression and chromatin accessibility programs upon entry to the T cell development and 2) essential TFs during the initiation of T-lineage program while T cell precursors are transitioning from the pre-thymic bone marrow progenitor states. To achieve this goal, a recently developed *in vitro* HSPC expansion technique was adapted to overcome the scarcity of progenitor populations. First, we thoroughly examined the T cell potential of expanded HSPCs under *in vivo* and *in vitro* environments with or without retroviral vector introduction to confirm their ability to generate T cells. Then, the molecular events during the launch of T cell programs were examined by comparing transcriptional profiles (measured by bulk and scRNA-seq) and chromatin accessibility features (determined by ATAC-seq) of expanded HSPCs and their progeny T cell precursors. To identify critical TFs in early T-development steps, 23 hematopoietic TFs were disrupted using the CRISPR-Cas9 system, which revealed novel roles for Lmo2, Erg, Hoxa9, and Meis1. Finally, the impacts of *Lmo2* knockout on the T cell program were defined by tracking their T cell development rate and gene expression changes.

### Animals and cell lines

C57BL/6J (B6. WT, Jax #000664), C57BL/6J-*Ptprc*^em6Lutzy^/J (B6. CD45.2, Jax #033076), B6.Cg-Tg(BCL2)25Wehi/J (Bcl2-tg, Jax #002320), B6.Gt(ROSA)26Sortm1.1(CAG-cas9*,-EGFP)Fezh/J (Cas9, Jax #026179) mice were purchased from the Jackson Laboratory and bred at the California Institute of Technology (Caltech). B6.Bcl11b^mCitrine/mCitrine^ (B6.Bcl11b-mCitrine reporter) mice were described previously (*38*). Both male and female mice were used for this study. All animals were bred and maintained under specific pathogen-free conditions at Caltech according to Institutional Animal Care and Use Committee (IACUC) regulations. For experiments involving chimeras, the busulfan priming and cell transfer followed a published protocol (*70*). For details, see Supplemental Methods.

### In vitro HSPC expansion culture

*In vitro* HSPC expansion was carried out under minor modifications of previously published conditions (*25, 71*), using bone marrow from 8-12 week-old B6 mice as input. The bone marrow populations were first sorted to isolate Lin^-^, Kit^+^, Sca-1^+^, EPCR^+^ cells for input to the cultures, and expanded HSPCs were periodically re-sorted during the 2-4 weeks of culture or enriched using magnetic beads to expand LSK cells selectively. Where noted, Flt3L (10 ng/ml) was added to the cultures for 4-5 days. For promoting T-cell development, the OP9-Dll1 (*36*) and MS5-mDll4 (*34, 35*) stromal cell lines were previously described, and both monolayer cocultures and ATO cultures to promote T-cell development followed our previously published procedures (*51*). Detailed conditions for cell purification, antibodies used, and culture variants are given in Supplemental Methods.

### Bulk and single-cell RNA-seq

Total RNA was isolated from 50,000 ∼ 300,000 sorted expanded HSPCs or sorted pro-T cells using the RNeasy Mini Kit (Qiagen #74105) with on-column DNase I treatment (Qiagen #1023460). Single-cell RNA-seq was then performed as described (*37*) using hashtag oligo labeling (BioLegend TotalseqA) to distinguish cells from different samples. Single-cell RNA-seq cDNA libraries were prepared using the 10X Chromium 3’ capture v3.1 kit and sequenced to a depth of 70,000-80,000 reads/cell for mRNA, 2000-2500 reads/cell for hashtags. Details are given in Supplemental Methods.

### ATAC-seq and CUT&RUN analyses

Samples for ATAC-seq were prepared according to a modification of the protocol for nuclear isolation described in ref. (*72*), with details given in Supplemental Methods. ATAC libraries were prepared using a published protocol (*73*) and sequenced to a targeted depth of 20 million reads per sample. CUT&RUN detection of activating and repressive histone modifications (H3K27ac and H3K27me3, respectively) was carried out following the original methods (*74–76*) with minor modifications (*51*). Libraries were prepared following a published method (*77*), and samples were sequenced to a depth of 10 million reads per sample. For details, see Supplemental Methods.

### Genomic analysis methods

Comprehensive details of all the analyses used here for bulk RNA-seq, scRNA-seq, ATAC-seq and CUT&RUN are provided in the Supplemental Methods.

### Statistical Methods

All conclusions were based on a minimum of two independent experiments (genomic analyses) and on 3-10 independent experiments (cell biological analyses). Details of the statistical tests used for every figure and supplemental figure in this paper are given in Supplemental Methods.

## List of Supplementary Materials

Supplementary text:

Supplemental Materials and Methods

Supplementary Figure Legends

Supplementary Table Legends

Supplementary Figures (Fig. S1 – Fig. S7)

Supplementary Tables (Table S1 – Table S5)

## Acknowledgments

We gratefully acknowledge Adam Wilkinson and members of his group (Oxford) for generous advice and encouragement; Kenneth Dorshkind and Encarnacion Montecino-Rodriguez (UCLA) for recommending the busulfan conditioning protocol for chimeras; Andrew S.-H. Koh (University of Chicago) for advice on ATAC protocol optimization, and members of the Rothenberg group for helpful discussions. OP9-Dll1 cells and MS5-mDll4 cells were originally the kind gifts of J. C. Zuñiga-Pflücker (University of Toronto and Sunnybrook Research Institute) and Gay Crooks (UCLA), respectively. We also thank Igor Antoshechkin and Vijaya Rao of the Muriel and Millard Jacobs Genetics and Genomics Laboratory, Caltech, for sequencing; Diane Trout and Henry Amrhein for sequence analysis system administration; Rochelle Diamond, Michael Gregory, Diana Perez, Jamie Tijerina, Madeleine Adolf, and Olivia Finney of the Flow Cytometry and Cell Sorting Facility, Caltech, and Lucy Brown, Ni Feng, and Shaun Hsueh of the City of Hope Analytical Cytometry Core for flow cytometry support; and Ingrid Soto (Office of Laboratory Animal Research), Maria Quiloan, and Mei Chau for care and supervision of the mouse colony.

## Authors contribution

B.S., S.J.C., and E.V.R. designed the project and wrote the manuscript. B.S., S.J.C., B.W.M, T.S., and B.A.W. conducted the experiments. B.S., S.J.C., and B.W.M. analyzed data. E.V.R. supervised research and acquired funding. All authors edited the paper and provided helpful comments.

## Accession numbers

All sequencing data have been deposited in Gene Expression Omnibus (GEO) under accession numbers GSE286217, GSE287757 (*in vivo* thymocyte bulk ATAC-seq), and GSE287751 (*Lmo2* knockout bulk RNA-seq).

## SUPPLEMENTAL MATERIALS AND METHODS

### *In vitro* hematopoietic stem and progenitor cell (HSPC) expansion

*In vitro* HSPC expansion was performed by following previous publications (*25, 71*). First, bone marrow was obtained from the femurs and tibiae of 8-12 week-old B6. WT or B6.Bcl11b^mCitrine/mCitrine^, or progeny of B6.Cas9;Bcl2 mice. The mature lineage^+^ cells were depleted using biotinylated antibodies against TCRβ (clone H57-597 BioLegend #109203), TCRγδ (clone GL-3, eBioscience #13-5711-82), CD3ɛ (clone 145-2C11, BioLegend # 100304), CD19 (clone 1D3, BioLegend #115503), B220 (clone RA3-6B2, eBioscience #13-0452-85), NK1.1 (clone PK136, BioLegend #108704), CD49b (clone HMa2, BioLegend #103522), CD11b (clone M1/70, BioLegend #101203). CD11c (clone N418, BioLegend #117303), Ly6G/C (clone RB6-8C5, BioLegend #108403), and Ter119 (clone TER-119, eBioscience #13-5921-82), followed by incubation with streptavidin microbeads (Miltenyi Biotec # 130-048-101) and magnetic separation using MACS LS columns (Miltenyi Biotec #130-042-401) by following the manufacturer’s protocol. Then, live (viability dye^-^) Lineage^-^ (Lin^-^) Kit^+^ (clone 2B8) Sca1^+^ (clone D7) CD150^+^ (clone TC15-12F12.2) and EPCR^+^ (clone RCR-16) cells were sorted. Sorted HSPCs were cultured at 200-1000 cells per well in a 48-well CellBIND plate (Corning #3338) in 500 μl of HSPC expansion medium (F12 DMEM (Gibco #11320-033), 1 mg/mL of PVA (Millipore Sigma #P8136), 1× Insulin–transferrin–selenium–ethanolamine (Gibco #51500-056), 10 mM HEPES (Gibco #15630-080), 1× Penicillin–streptomycin–glutamine (PSQ, Gibco #0378-016); filtered using a 0.22 μm filter) freshly supplemented with 100 ng/mL of recombinant mouse thrombopoietin (TPO, PeproTech #AF-315-14 or BioLegend #593304) and 10 ng/mL of recombinant mouse stem cell factor (SCF, PeproTech # 250-03) at 37°C, 5% CO_2_ environment. After the initial 5 days of culture, complete media change was performed every 2-3 days. Before the culture reached 80% confluency, cells were split into 2-3 wells or one well of a 24-well CellBIND plate (Corning #3337). At days 14-28, expanded HSPCs were further purified by positively selecting the Sca1^+^ cells using MACS MS columns (Miltenyi Biotec #130-042-201) or sorted for Lineage^-^Kit^+^Sca1^+^(LSK) or CD150^+^EPCR^+^ LSK phenotype.

### Retroviral transduction

Non-tissue culture plates (Corning #351147) were coated with 50 μg/mL RetroNectin (Takara bio #T100B) 1×PBS at 4°C overnight. After the removal of unbound RetroNectin, retroviral supernatant was added to the plate and centrifuged at 2000×g, 32°C for 2 hours. Freshly isolated bone marrow progenitors or expanded progenitors were then incubated with the retroviral vector-bound plate at 37°C overnight in a 5% CO_2_ incubator. To ensure progenitors were in cell cycle, fresh progenitors were infected in OP9 media (αMEM (Gibco #41061-029), 20% FBS, 1×PSQ, and 50 μM β-mercaptoethanol (Gibco #21985-023)) supplemented with 10 ng/mL of recombinant human IL-7 (PeproTech #200-07), 10 ng/mL of recombinant human Flt3-ligand (Flt3L, PeproTech #300-19), and 10 ng/mL of SCF; and expanded HSPCs were infected in HSPC expansion media with 100 ng/mL TPO and 10 ng/mL SCF. For Flt3L-primed expanded HSPCs, 10 ng/mL of Flt3L was additionally provided. After infection, cells were removed from the viral particles and cytokines, and subsequently sorted and transferred into mice or co-cultured with OP9-Dll1 cells.

### Bone marrow chimera reconstitution

Bone marrow chimera experiments were performed by following a previously published protocol (*70*). Briefly, busulfan (Millipore Sigma #B2636-10g, stock: 15 mg/mL in DMSO) was diluted in 37 °C 1× PBS to the 1.5 mg/mL working concentration and sterilely filtered using a 0.2 μm nylon filter before the injection. Eight-to-12-week-old CD45.1^+^CD45.2^+^ hosts (F_1_ progeny of B6. CD45.1 and CD45.2 mice) were treated with 25 mg/kg body weight of busulfan by intraperitoneal injection on days −2 and −1 prior to bone marrow transplantation. One day after the second dosage of busulfan treatment (on day 0), 4-5×10^5^ of freshly isolated Lin^-^ bone marrow progenitor cells or expanded HSPCs from sex-matched CD45.1^-^CD45.2^+^ mice were transferred via retro-orbital injection. To test T cell competence in retrovirally-transduced progenitors, freshly isolated Lin^-^ bone marrow progenitors or expanded HSPCs were infected with MSCV-mCherry empty vector (Addgene #80157) or E42-mTurquoise2 control vector (Addgene #189799) as described above. After an overnight infection, infection^+^ (either mCherry^+^ or mTurquoise2^+^) Lin^-^ cells were sorted by flow cytometry and transferred into busulfan-conditioned recipients (1×10^5^ cells per mouse). To evaluate the reconstitution rate and T cell potentials, the thymus and the spleen of the recipient animals were analyzed at 4-6 weeks post-transfer. As busulfan is a chemotherapeutic agent, it was handled following the Caltech guidelines for the use of cytotoxic or chemotherapeutic drugs.

### *In vitro* artificial thymic organoid culture (ATO) and OP9-Dll1 co-culture

To test T-lineage potential and monitor fine-scale time course events of T cell development, freshly isolated bone marrow progenitor cells or *in vitro*-expanded HSPCs were compared in an ATO 3-dimensional culture system by following the original method. Briefly, 1,000 progenitor cells and 150,000 mouse MS4-Dll4 cells (originally obtained from Dr. Gay Crooks (*34, 35, 78*)) were aggregated and seated on a culture insert (Millipore Sigma #PICM0RG50) in serum-free ATO medium (DMEM-F12, 1× B27 (Gibco #17504-044), 1× PSQ, and 30 μM Ascorbic acid (Millipore Sigma # A8960, freshly reconstituted in 1×PBS at 30 mM)) supplemented with 5 ng/mL of IL-7 and 5 ng/mL of Flt3L. The cytokine-supplemented culture medium was replaced every 3 days and IL-7 and Flt3L concentrations were dropped to 1 ng/mL (each) after day 10. ATO cultures were done under 37 °C, 5% CO_2_ environment.

To track early T cell development, freshly isolated bone marrow progenitor cells or *in vitro*-expanded HSPCs were co-cultured with OP9-Dll1 cells (originally obtained from Dr. J. C. Zúñiga-Pflücker (*36*)) with 1o ng/mL of IL-7 and 10 ng/mL of Flt3L (from days 0-5) or 5 ng/mL of IL-7 and 5 ng/mL of Flt3L (from days 6-8) or 1 ng/mL of IL-7 and 1 ng/mL of Flt3L (from days 8 to later) in OP9 medium. Progenitor cells co-cultured with OP9-Dll1 stroma were split and replated on new OP9-Dll1 cells every 3-4 days with a complete medium change. OP9-Dll1 *in vitro* co-cultures were incubated in a 37 °C, 7% CO_2_ environment.

### Isolation of *in vivo* DN thymocytes

Thymi were harvested from 4.5-7-week-old mice. 8-10 sex-matched thymi were pooled for each ATAC replicate. At least one ATAC replicate per DN population was made from male and female cells, each. Whole thymi were manually dissociated using 40-micron cell strainers (Falcon cat# 352340). CD3-expressing cells were then depleted using PNA panning at a density of 5×10^7^ cells per 10-cm petri dish for 70 minutes at 4°C with occasional swirling. Following PNA panning, cells were again filtered through a 40-micron cell strainer, and Fc receptors were blocked by incubating the cells in 2.4G2 hybridoma cell (ATCC #HB-197) supernatant for 10-15 minutes on ice at a density of 10^8^ cells per mL. Cells were then incubated for 20 min on ice in CBH buffer (1×HBSS, 0.5% BSA, and 10 mM HEPES) at the same concentration containing a biotin-conjugated lineage cocktail: CD3 (1:100), CD4 (1:100), CD8α (1:100), TCRβ (1:100), TCRγδ (1:100), CD19 (1:200), NK1.1 (1:200), CD49b (1:200), CD122 (1:200), CD11b (1:200), CD11c (1:200), Ter119 (1:100), and Ly6G/C (1:100). Cells were then washed and incubated for 10 min on ice in CBH containing 1:10 Streptavidin microbeads (Miltenyi) on ice at 10^8^ cells per mL and subsequently depleted by magnetic separation using MACS LS columns, 1 mL cells per column, according to the manufacturer’s protocol. The lineage-depleted cells were then stained with fluorescent antibodies targeting CD45, CD44, Kit, and CD25, as well as fluorescent-labeled streptavidin as described above. Cell viability was determined using a near-IR fixable viability dye (Invitrogen #L10119) according to the manufacturer’s protocol. Cells were sorted as described below. ATAC-seq was performed on 20,000-30,000 cells for ETP, ETP-trans, and DN2a samples; 30,000-50,000 cells for DN2b samples, based on the overall cell recovery, as described later.

### Flow cytometry analysis and cell sorting

Cell surface staining was performed following Fc blocking by incubating single cell suspensions in 2.4G2 hybridoma cell supernatant. Then cells were stained with a biotin-conjugated lineage cocktail: TCRβ, TCRγδ, CD19, NK1.1, CD49b, CD11b, CD11c, and Ly6G/C. Secondary surface staining was performed with fluorescently conjugated streptavidin and antibodies (listed below) in CBH buffer. A viability dye (Life Technologies, Aqua) or 7AAD (eBioscience) was applied by following manufacturers’ protocols to exclude dead cells. Samples were acquired using a CytoFlex analyzer (Beckman Coulter), and data was analyzed with FlowJo v.10.10.0 (BD). Cells were gated on live, singlets, mature lineage (TCRβ, TCRγδ, CD19, NK1.1, CD49b, CD11b, CD11c, Ly6G/C)^-^ (Lin^-^) population for sorting and analysis.

Cell sorting was performed by following the same staining protocol described above. BD FACSAria Fusion and CytoFlex SRT (Beckman Coulter) at Caltech Flow Cytometry Facility or BD FACSAria Fusion at City of Hope Flow Cytometry Facility were utilized for cell sorting.

Specifically, the following cell populations were sorted for each experiment.

1. “ELSK” (Live Lin^-^ Sca1^+^ Kit^+^EPCR^+^CD150^+^):

Input population of in vitro HSPC expansion in Figures 1-6.

Re-purified expanded HSPCs for bulk RNA-seq (Figure 2), scRNA-seq (Figure 3), ATAC-seq (Figure 4), and cytokine/chemokine treatment (Figure 5).

1. 2) “LSK” (Live Lin^-^ Sca1^+^ Kit^+^):

Re-purified expanded HSPCs for OP9-Dll1 co-culture and ATO culture (Figures 1, 2), bulk RNAseq (Figure 2), and CRISPR-Cas9 screening (Figure 6). Also, fresh bone marrow LSKs were sorted for *in vitro* OP9-Dll1 in vitro culture (Figure 2, Figure 6) and scRNA-seq (Figure 3).

1. 3) “ETP” (Early T-progenitors, a subset of DN1): Gated on Live Lin^-^ CD45^+^ Kit^+^ CD25^-(^Figures 2, 5, and 6).
2. 4) “DN1”: Gated on Live Lin^-^ CD45^+^ CD44^+^ CD25^-^ (Figure 7).
3. 5) “Transitional ETP”: Gated on Live Lin^-^ CD45^+^ Kit^+^ CD25^lo^ (Figure 4, bulk thymic ATAC).
4. 6) “DN2”: Live Lin^-^ CD45^+^ Kit^+^ CD25^+^ (Figures 2, 5, 6) or Live Lin^-^ CD45^+^ CD44^+^ CD25^+^Thy1^+^ (Figure 7).
5. 7) “Bcl11b^-^ DN2a”: Live Lin^-^ CD45^+^ Kit^+^ CD25^+^ Bcl11b-mCitrine^-^ (Figures 2, 5).
6. 8) “DN2a”: Live Lin^-^ CD45^+^ CD44^+^ Kit^high^ CD25^+^ (Figure 4, bulk ATAC thymic DN2a).
7. 9) “Bcl11b^+^ DN2a/b”: Live Lin^-^ CD45^+^ Kit^high-int^ CD25^+^ Bcl11b-mCitrine^+^ (Figures 2, 5).
8. 10) “DN2b”: Live Lin^-^ CD44^+^ Kit^int^ CD25^+^ (Figure 4, bulk ATAC thymic DN2b).

The following antibodies were used for flow cytometry analysis and cell sorting:

| Marker | Conjugate | Clone | Manufacturer | Cat # |
| --- | --- | --- | --- | --- |
| B220 | Biotin | RA3-6B2 | eBioscience | 13-0452-85 |
| CD11b | Biotin | M1/70 | BioLegend | 101204 |
| CD11c | Biotin | N418 | BioLegend | 117304 |
| CD150 | PEcy7 | TC15-12F12.2 | BioLegend | 115913 |
| CD19 | Biotin | 6D5 | BioLegend | 115504 |
| CD25 | AF647 | PC61 | BioLegend | 102020 |
| CD25 | APC | PC61.5 | Invitrogen | 17-0251-82 |
| CD25 | e450 | PC61.5 | Invitrogen | 48-0251-82 |
| CD25 | PE | PC61.5 | Invitrogen | 12-0251-83 |
| CD3ε | Biotin | 145-2C11 | BioLegend | 100304 |
| CD4 | BV510 | RM4-5 | BD Horizon | 563106 |
| CD4 | PE | RM4-5 | BioLegend | 100512 |
| CD44 | e450 | IM7 | Invitrogen | 48-0441-82 |
| CD44 | PE | IM7 | BioLegend | 103008 |
| CD44 | PerCPcy5.5 | IM7 | BioLegend | 103032 |
| CD44 | APC | IM7 | BioLegend | 103012 |
| CD45 | AF488 | 30-F11 | BioLegend | 103122 |
| CD45 | PEcy7 | 30-F11 | Invitrogen | 25-0451-82 |
| CD45.1 | PEcy7 | A20 | BioLegend | 110730 |
| CD45.2 | APC | 104 | BioLegend | 109813 |
| CD45.2 | APCe780 | 104 | Invitrogen | 56-0454-81 |
| CD45.2 | APCe780 | 104 | eBioscience | 47-0454-82 |
| CD49b | Biotin | DX5 | Invitrogen | 13-5971-85 |
| CD8α | APC | 53-6.7 | Invitrogen | 17-0081-82 |
| CD8α | APCe780 | 53-6.7 | Invitrogen | 47-0081-82 |
| CD8α | Biotin | 53-6.7 | BioLegend | 100704 |
| Kit | e450 | 2B8 | eBioscience | 48-1171-82 |
| Kit | PE | 2B8 | Invitrogen | 12-1171-83 |
| EPCR | APC | 1560 | Invitrogen | 17-2012-82 |
| EPCR | APC | RCR-16 | BioLegend | 141506 |
| hNGFR | APC | ME20.4 | BioLegend | 345108 |
| hNGFR | SB436 | ME20.4 | Invitrogen | 62-9400-42 |
| Ly6G/C | Biotin | RB6-8C5 | Invitrogen | 13-5931-86 |
| NK1.1 | Biotin | PK136 | BioLegend | 108704 |
| Sca1 | Biotin |  | BioLegend | 108103 |
| Sca1 | PE | D7 | eBioscience | 12-5981-83 |
| Streptavidin | APCe780 |  | eBioscience | 47-4317-82 |
| Streptavidin | BV510 |  | BioLegend | 405233 |
| Streptavidin | PerCPcy5.5 |  | BioLegend | 405214 |
| TCRβ | Biotin | H57-597 | BioLegend | 109204 |
| TCRβ | e450 | H57-597 | eBioscience | 48-5961-82 |
| TCRβ | PE | H57-597 | eBioscience | 12-5961-83 |
| TCR$\gamma\delta$ | Biotin | GL-3 | Invitrogen | 13-5711-85 |
| TCR$\gamma\delta$ | e450 | GL-3 | Invitrogen | 48-5711-82 |
| TCR$\gamma\delta$ | PEcy7 | GL-3 | BioLegend | 118124 |
| Ter119 | Biotin | Ter119 | Invitrogen | MA5-17820 |
| Thy1<br>(CD90.2) | APCe780 | 53-2.1 | Invitrogen | 47-0902-82 |
| Thy1<br>(CD90.2) | FITC | 30-H12 | Invitrogen | 11-0903-82 |

### Limiting dilution analysis

To estimate progenitor frequency for the T-lineage, expanded HSPCs were sorted for EPCR^+^CD150^+^LSK phenotype and subjected to 4 days of unprimed condition (the same expansion condition) or Flt3L-priming condition (expansion condition + 10 ng/mL Flt3L). Then, cells were washed with OP9 medium twice and subjected to the OP9-Dll1 co-culture. OP9-Dll1 cells were prepared one day prior to the start of T cell differentiation by seeding 5000 cells per well in 96-well plates. Limiting dilution was performed by plating 1000, 100, 30, 10, 3, or 1 cell per well, and each condition had 3-24 wells. After 5 days, surviving CD45^+^ Lin^-^ cells were counted using PE beads (BD Biosciences #340495).

A well with the day 5 cell number (live CD45^+^ lineage^-^) > the d0 input cell number was considered a success. The progenitor frequency and the significance between unprimed and Flt3L-primed conditions were calculated using the Extreme Limiting Dilution Analysis (ELDA) tool (*79*).

### Bulk RNA-seq

Total RNA was isolated from 50,000 ∼ 300,000 sorted expanded HSPCs or pro-T cells using the RNeasy Mini Kit (Qiagen #74105) with on-column DNase I treatment (Qiagen #1023460) by following the manufacturer’s protocols. Sequencing libraries were constructed at the Caltech Genomics Facility and sequenced on Illumina NextSeq (1×50 base pair, targeted to paired-read, 30 million read depths per sample).

### Single-cell RNA-seq

Single-cell RNA-seq was performed as described (*37*). On days 0, 5, and 9 after co-culture with OP9-Dll1 cells, progenitor cells were stained with biotinylated antibodies as described above. During the secondary surface antibody staining step, different unique hashtag-oligo (HTO) antibodies (TotalseqA HTO1-HTO11, clones M1/42; 30-F11, BioLegend #155801, #155803, #155805, #155807, #155809, #155811, #155813, #155815, #155817, #155819, #155821) were added to distinguish each sample, then cells were sorted for indicated markers using the BD FACSAria Fusion at the Caltech Flow Cytometry Facility. After FACS sorting the target cells, samples were washed with 1×HBSS supplemented with 0.5% BSA and 10 mM HEPES and resuspended to 1×10^6^ cells/1 mL concentration. Then, 1.6 – 2.0 x10^5^ cells were loaded into a 10X Chromium lane, and the subsequent preparation was conducted following the instruction manual of 10X Chromium v3.1. Single-cell RNA-seq cDNA libraries were prepared using 10X Chromium 3’ capture v3.1 kit. Single-cell hashtag oligo library was prepared in accordance with the BioLegend TotalseqA guide. Constructed single-cell RNA-seq cDNA libraries were sequenced to a depth of 70-80,000 reads per cell and hashtag oligo libraries were sequenced for 2,000-2,500 reads per cell.

The following conditions were used for each scRNA-seq experiment. Experiment 1:

HTO 1: freshly isolated EPCR^+^ CD150^+^ LSK co-cultured with OP9-Dll1 for 9 days

HTO 2: EPCR^+^ CD150^+^ LSK sorted expanded HSPC co-cultured with OP9-Dll1 for 9 days

HTO 3: freshly isolated EPCR^+^ CD150^+^ LSK co-cultured with OP9-Dll1 for 5 days

HTO 4: EPCR^+^ CD150^+^ LSK-sorted expanded HSPC co-cultured with OP9-Dll1 for 5 days

HTO 5: LSK-sorted Flt3L-primed expanded HSPC co-cultured with OP9-Dll1 for 5 days

HTO 6: freshly isolated EPCR^+^ CD150^+^ LSK

HTO 7: expanded HSPC EPCR^+^ CD150^+^ LSK

HTO 8: Flt3L-primed expanded HSPC LSK

Experiment 2:

HTO 1: freshly isolated EPCR^+^ CD150^+^ LSK

HTO 2: freshly isolated LSK

HTO 3: LSK-sorted Flt3L-primed expanded HSPC

HTO 4: LSK-sorted expanded HSPC

HTO 5: freshly isolated EPCR^+^ CD150^+^ LSK co-cultured with OP9-Dll1 for 5 days

HTO 6: LSK-sorted expanded HSPC co-cultured with OP9-Dll1 for 5 days

HTO 7: Flt3L-primed expanded HSPC LSK co-cultured with OP9-Dll1 for 5 days

HTO 9: freshly isolated EPCR^+^ CD150^+^ LSK co-cultured with OP9-Dll1 for 9 days

HTO 10: LSK-sorted expanded HSPC co-cultured with OP9-Dll1 for 9 days

HTO 11: Flt3L-primed expanded HSPC LSK co-cultured with OP9-Dll1 for 9 days

Note: Figures S5E, F only show the results from experiment 2 to directly compare Flt3L-priming effects within the same experiment.

### ATAC-seq

Nuclei of progenitor cells were isolated by incubating cells in 0.04 % NP-40 and 0.1 % Tween-20 in detergent-free ATAC lysis buffer (130 mM NaCl, 5 mM MgCl_2_, 5 mM Kcl, 0.05 mM EDTA, 10% v/v glycerol, and 25 mM HEPES; protocol is a modification of the ATAC lysis buffer employed by Gamble et al for nuclei isolation from DN thymocytes (*72*)) for 1 minute on ice at 1,000 cells/μL concentration (for expanded HSPCs and their progeny DNs) or 0.1% NP-40 and 0.1% Tween-20 in detergent-free ATAC lysis buffer for 2 minutes on ice at 1,000 cells/μL concentration (for adult *in vivo* thymic progenitors). After incubations, cells were immediately washed with 500 μL of detergent-free ATAC lysis buffer and centrifuged at 400 × *g* for 5 minutes at 4 °C. Then, nuclei were resuspended in the transposition mix (1× TD buffer (Illumina #15027866), 33% PBS, 0.01% digitonin, 0.1% Tween-20, and Tn5 transposase (Illumina #15027916) in UltraPure H_2_O (Invitrogen #10977-015)) at 1,000 cells/μL concentration and incubated at 37 °C for 30 minutes in a thermocycler. Following transposition, DNA purification was immediately performed using a DNA clean and concentrator kit (Zymo #D4014) according to the manufacturer’s protocol. ATAC libraries were constructed by following a previously published protocol (*73*). Briefly, DNA was again purified using the Zymo DNA clean and concentrator kit following PCR amplification according to the manufacturer’s protocol. Small nucleosome-free, 1-, and 2-nucleosome fragments were then enriched using double-sided size selection (0.6X SPRI and 1.8 X SPRI) using AMPure XP beads (Beckman Coulter #A63881). DNA concentration was assessed by Qubit HD DNA assay (Invitrogen cat# Q32854), and molarity was calculated by Bioanalyzer HS DNA kit (Agilent cat# NC1738319). After the library preparation, the sequencing was performed with paired-end sequencing of 1×50 base pair, a targeted depth of 20 million reads per sample on Illumina NextSeq at the Caltech Genomics Facility.

### CUT&RUN (C&R)

C&R was performed by following the original methods previously described (*74–76*) with minor modifications as previously reported (*51*). Briefly, untreated or Flt3L-primed expanded HSPCs were washed with wash buffer (100 mM NaCl, 20 mM HEPES, 0.5 mM spermidine, 1X protease inhibitor, and 0.5% BSA) and 2-2.5×10^5^ cells were bound to 10 μL of activated concanavalin-A coated beads by incubating in wash buffer at room temperature for 5-10 min. The bead-bound cells were incubated with anti-rabbit antibodies for H3K27ac (Abcam #ab4729), H3K27me3 (Cell Signaling Technology #9733), or negative control antibody (guinea pig anti-rabbit antibody, Antibodies-Online #ABIN101961) by incubating in 200 μL of antibody buffer (0.0005-0.001% wt/vol digitonin in wash buffer with 1 mM EGTA, 1-2 μg antibody) for 2 hours at 4 °C. After antibody incubation, permeabilized cells were washed with digitonin buffer (0.0005-0.001% wt/vol digitonin in wash buffer) and incubated with 700 ng/mL of protein A-MNase (pA-MN, purified from Addgene #86973 by Caltech Protein Expression Center) in a total volume of 250 μL for 1 hour at 4 °C. For chromatin digestion, cells were washed with low-salt rinse buffer (0.5 mM spermidine, 20 mM HEPES, 0.0005-0.001% digitonin, 1x protease inhibitor) and incubated with 200 μL of low-salt high-Ca^2+^ incubation buffer (3.5 mM HEPES, 10 mM CaCl_2_, 0.0005-0.001% digitonin) at 0 °C for 5 min. After digestion, the incubation buffer was quickly replaced with 200 μL of 1× STOP buffer (170 mM NaCl, 20 mM EGTA, 50 μg/mL RNAse, 25 μg/mL glycogen). Digested chromatin was released by incubating at 37 °C for 15 min and centrifuged at 4 °C at 16,000g for 5 min. DNA was extracted by incubating with 0.1% SDS and 250 μg/mL of Proteinase K at 50 °C for 1 hour, followed by Phenol Chloroform extraction.

C&R libraries were prepared using the NEBNext ChIP-Seq Library Preparation Kit (NEB #E7645S, #E7500S, #M5505S) by following a previously published protocol (*77*). After the library preparation, the sequencing was performed with paired-end sequencing of 1×50 base pair, a targeted depth of 10 million reads per sample on Illumina NextSeq at the Caltech Genomics Facility.

### Bulk RNA-seq analysis

The sequencing alignment and gene expression calculation were performed by following the ENCODE RNA-seq pipelines. Briefly, the adapter trimmed reads were aligned to the mouse reference genome GRCm38/mm10 using STAR (v 2.5.1b), and gene expression was calculated with RSEM (v 1.2.31). Then, significant changes in gene expression were determined using DEseq2 (v 1.46.0). For the DEseq2 input matrix, low-count genes were filtered out by keeping at least one sample with greater than 8 counts or the sum of all samples was greater than 20.

Supervised Principal Component Analysis (PCA) was performed as described previously (*40*). Briefly, the PC loadings for 64 informative T-developmental trajectory genes were determined from a previous single-cell RNA-seq study (*18*), and the PC1 and PC2 loadings were projected to bulk RNA-seq data.

Differentially expressed genes (DEGs) were defined by average TPM (across compared samples) ≥ 5, adjusted *p-*value < 0.01, and absolute Log_2_ fold-change > 1 (i.e. at least twofold increased or decreased), whereas non-DEGs were defined by average TPM ≥ 5, adjusted *p-*value ≥ 0.01 or absolute Log_2_ fold-change < 1 for the results shown in Figures 2, 4, 5, and 6. To define DEGs in Fig. 7 and Fig. S7, a more permissive threshold was used, with adjusted *p*-value < 0.05 and |(Log_2_ fold-change)| > 0.5 for DEGs.

For creating heatmaps of Figures 2, S2, and S6, hierarchical clustering was performed using R hclust function (Manhattan distance, complete linkage) and gene expression data were visualized as *Z*-scores (calculated from TPM values) using R pheatmap.

Gene set enrichment analysis (GSEA) was performed by pre-ranking RNA-seq results according to a shrinkage of log_2_ fold changes calculated using DEseq2. The gene sets from the hallmark gene sets (H), curated gene sets (C2), regulatory target gene set (C3), computation gene set (C4), ontology gene sets (C5), and immunologic signature gene sets (C7) of the Molecular Signatures Database (MsigDB) were used for computing enrichment. The normalized enrichment score (NES), nominal p value, and false detection rate (FDR) q-value, and leading edge genes were determined using fGSEA (v 1.32.0). The significant pathways obtained by FDR q value < 0.05 were visualized using Python Matplotlib (v 3.7.2) and the leading edge genes overlapped with DEGs were plotted as heatmaps as described above.

### Single-cell RNA-seq analysis

The raw reads from cDNA libraries were aligned to the mouse reference genome GRCm38/mm10 using CellRanger (v 6.1.2) and the hashtag oligo (HTO) libraries were quantified and demultiplexed using in-house tools (hashtag tool) as described previously (*37*).

For QC and downstream analysis, Seurat (v 5.1.0) was utilized. Briefly, cells displaying unique hashtag oligo identity (to exclude doublets labeled by multiple HTOs) and expressing at least 500 genes were considered. Potential doublets expressing more than 7800 genes and unhealthy cells displaying high mitochondrial RNA contents (greater than 0.18 %) were further filtered out.

Two independent experiments and two different previously published scRNA-seq datasets from our group (*37, 51*) were integrated with the reciprocal principal component analysis (PCA) algorithm using the 16,000 integration features and 5 anchors. After data integration, UMAP was constructed using the first 30 PCs, and differential gene expression analysis was performed by using pseudocount 0.1. Differential expression test was performed by non-parametric Wilcoxon rank sum test using Seurat, and pseudo-bulk DEGs were determined by the genes expressed by a minimum 1% of cells, adjusted *p-* value < 0.01, and absolute fold-change ≥ 2 (Fig 3) or the genes expressed by a minimum 1% of cells, adjusted *p-*value < 0.05, and absolute fold-change ≥ 1.5 (Fig. 4) or the genes expressed by a minimum 1% of cells, adjusted *p-*value < 0.05, and absolute Log_2_fold-change > 0.5 (Fig. S5). The pseudotime inference was performed using Monocle (v 3.14), and the root principal node was defined based on LT-HSC features (*Mecom* > 3 & *Tcf15* > 1 & *Kit* > 1 & *Procr* > 1 & *Ly6a* > 1 & *Il7r* < 0.5 & *Flt3* < 0.5).

### ATAC-seq analysis and CUT&RUN analysis

Sequenced reads were mapped to the GRCm38/mm10 mouse reference genome using Bowtie 2 (v 3.5.1), after which PCR duplicates were removed using Picard (v 2.18.29). Low-quality reads and reads mapped to blacklisted regions and the mitochondrial genome were filtere using Samtools (v 1.9). Peak calling was conducted using Genrich (v 0.6) for ATAC-seq or SEACR (v 1.3) for CUT&RUN (C&R) histone peaks with a stringent mode. For UCSC Genome Browser visualization, Bigwig files were generated from the aligned bam file using deepTools (v 3.5.1, bamCoverage--binSize 20--normalizeUsing CPM).

To compare different sources of ATAC-seq results, publicly available ATAC-seq raw sequenced reads (GSE100738) (*22*) were downloaded and processed as described above. After the Genrich peak calling, peaks with an average count > 3 across all samples were further considered for the downstream analysis. Then, quantile normalization was performed after log_2_ transformation of count matrix +0.1 with n bins = 100 using a custom script. The quantile normalized matrix was used to perform principal component analysis with DEseq2 (v 1.46.0).

To compare *in vivo* vs. *in vitro* ATAC profiles or expanded HSPCs and their progeny ATAC features, ATAC-seq results generated from this paper were only utilized to minimize the technical variances. Differential accessibility (DA) sites were calculated after filtering out lowly assessable peaks (average count ≤ 3). The genomic regions with absolute fold-change > 1 and adj *p*-value < 0.01 in each comparison were defined as DA sites using DEseq2 by inputting raw counts matrix without quantile normalization.

The peak-centered heatmaps were generated using deepTools2 (v 3.5.1) in a 3000 bp region (ATAC seq) or 4000 bp region (C&R). The distinct ATAC-seq groups were determined based on pair-wise comparisons of DA results, which were then categorized by employing a multinomial logistic regression model with the Limited-memory Broyden-Fletcher-Goldfarb-Shanno (lbfgs) solver using Python sklearn (v 1.3.0) by training each categorical group with 5-6 different pairwise comparison results. Co-occurring or unique peaks of C&R histone marks were computed using the HOMER package (v 4. 11. 1, mergePeaks-venn), and each class was defined by Boolean logic.

Genes associated with ATAC peaks were annotated using GREAT (v.4.0.4) with proximal:5kb upstream, 1kb downstream, plus distal: up to 1000 kb mode. Motif enrichment was analyzed using the HOMER package (findMotifGenome.pl) using a 150 bp window and *de novo* results were reported.

### Specific gRNA design and gRNA sequences

The 19-mer or 20-mer sgRNAs against target transcription factors were designed by using the CHOPCHOP web tool (https://chopchop.rc.fas.harvard.edu/) (*80*) or using the guide RNA sequences (against *Spi1*, *Erg*, *Meis1*+*Hoxa9* dual gRNA, and non-targeting control) published previously (*37, 39*). The designed sgRNAs were inserted into the empty E42 control vectors (Addgene # 189799, # 189800) as described previously. To increase the deletion efficiency of the target gene, 2-3 different sgRNA expressing vectors were generated for each target gene, and each duo or trio was pooled for packaging in Phoenix-Eco cells using Fugene 6 Transfection Reagent (Promega #E2691) according to manufacturer’s protocols.

Sequences of sgRNAs are below:

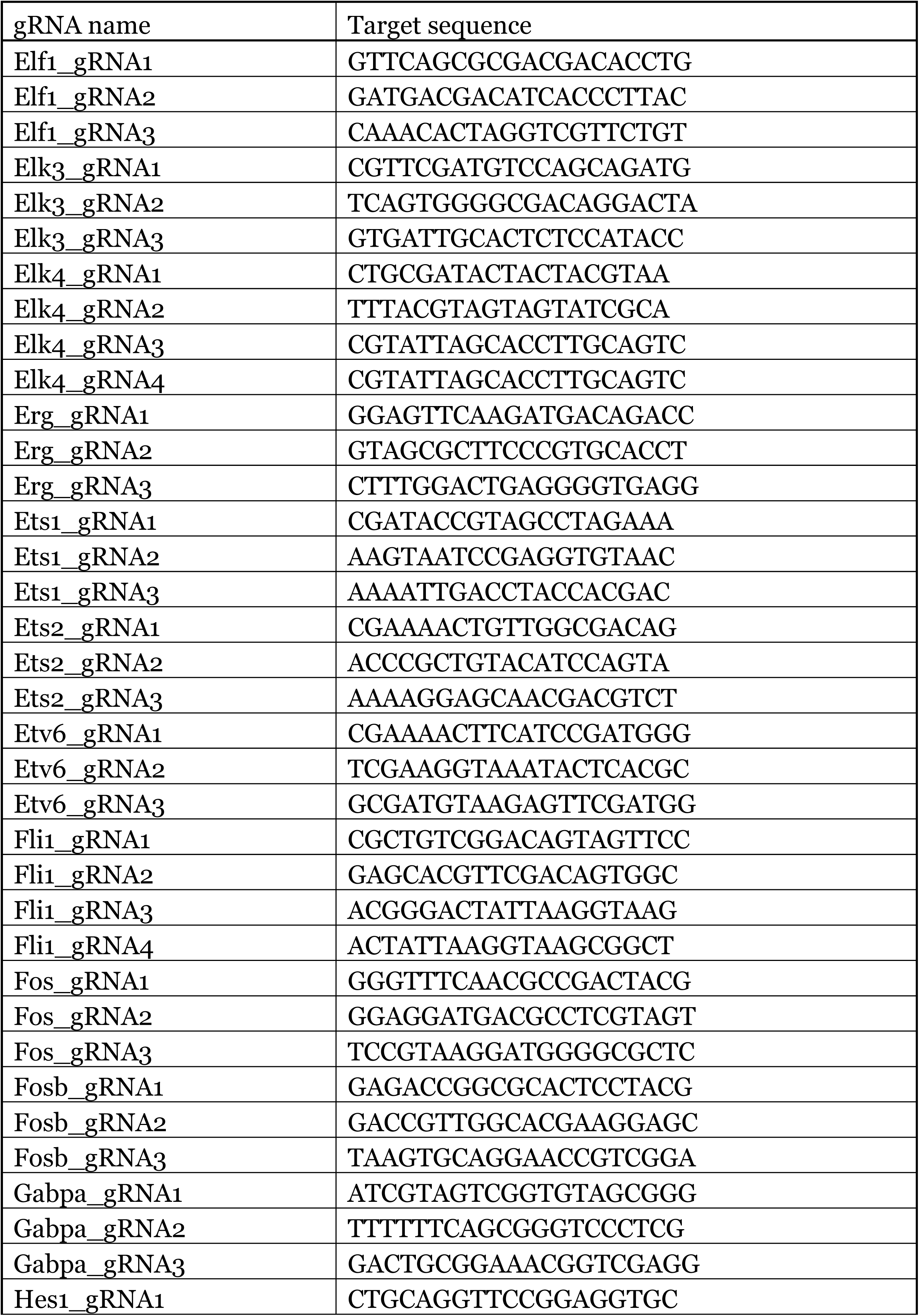
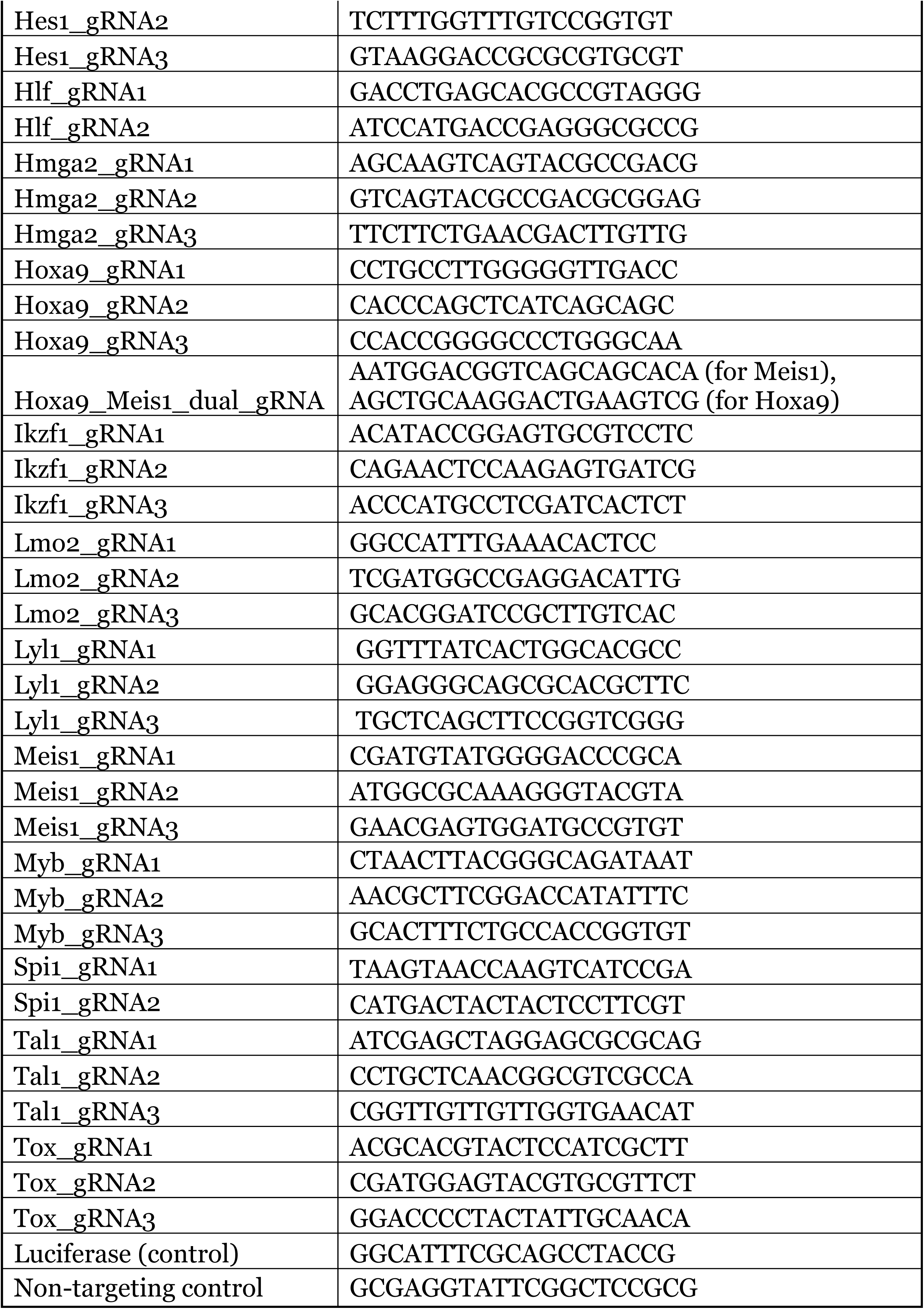

### *Lmo2* Knockout Experiments

These experiments used Lin^-^ progenitor cells from B6.Cas9; Bcl2 mice for HSPC expansion (Fig 7) and immediate infection (Fig S7B). Cells were collected and cultured as previously described in the HSPC expansion section, with slight alterations: 1) expanded HSPC cultures were not sorted but were stained at day 10-13 of expansion with Sca1-PE (1:50 dilution), followed by incubation with magnetic PE-beads to select for Sca1^+^ cells using MACS LS Columns and 2) Lin^-^ progenitors (Fig S7B) and Sca1^+^ expanded HSPCs (Fig 7) were frozen down in Bam Banker solution (FUJIFILM #302-14681) and then thawed for experimental use in the appropriate media conditions (Lin^-^ progenitors: overnight thaw in OP9 media with 10 ng/mL IL7, 10 ng/mL Flt3L, and 10 ng/mL SCF in a 5% CO_2_ environment, Sca1^+^ expanded HSPCs: thawed for 4-6 days in HSPC expansion conditions, or if Flt3L-primed, with an added 10 ng/mL Flt3L for 4 days of the thawing period). Infection with control vs. *Lmo2* KO vectors followed the previously described retroviral transduction section but used the corresponding thawing media rather than OP9 media for overnight incubation. Flow cytometry analysis and cell sorting methods are as previously described, with flow cytometry performed on the MACSQuant (Miltenyi) and analyzed with FlowJo. The panel for DN developmental progression used in these experiments includes CD44-PE (1:300), 7AAD (1:100), SA-PerCP-cy5.5 (1:300), CD45-PEcy7 (1:1000), CD25-APC (1:300), Thy1-APCe780 (1:300), and mTurquoise2 as the infection marker.

Bulk RNA-seq samples from day 5 co-cultures were prepared and analyzed as previously described; however, DEGs were defined by adjusted *p*-value < 0.05 and absolute Log_2_ fold-change > 0.5 (results shown in Fig 7E, H, I, raw data in Table 5). The expanded HSPC to Bcl11b^+^DN2a/2b DEGs shown in Fig 7E were defined by adjusted *p*-value < 0.05 and absolute Log_2_ fold-change > 0.8 using the bulk RNA-seq samples described in Fig 2 E-H. The heatmap in Fig 7G was generated with Seaborn’s clustermap using *Z*-scores across every gene, such that each gene has a mean of 0 and a variance of 1 across all conditions (calculated from TPM values).

Published DEG sets were taken from Table S1 from (*81*), where TF-inhibited and TF-activated targets were defined by published datasets:

Erg target genes (KO, scRNAseq): GSE165835 (*37*) Notch target genes (KO, bulk RNAseq): GSE148441 (*40*)

PU.1 target genes: GSE165835 (KO, scRNAseq) (*37*); GSE93755 (KO and gain of function, bulk RNAseq) (*82*)

TCF1 target genes: GSE165835 (KO, scRNAseq) (*37*); GSE33513 (KO, microarray)

(*83*); GSE26560 (KO, microarray) (*84*)

### Statistical analysis

The following statistical analysis was performed to compare the means of two groups or more than two groups using Prism software (v.10.4.1, GraphPad). Statistical comparisons of distributions and linear correlation were performed using scipy.stats. (v.1.9.1) using Python (v.3.8.13). ns = not significant, * = *p-*value < 0.05, ** = *p-*value < 0.01, *** = *p-*value < 0.001. Graphs with multiple data points were shown with median values and interquartile ranges. Tests used in individual figures were as follows.

**Figure 1:** (E, G) Unpaired t-test and (I) unpaired t-test with Welch’s correction for unequal standard deviation. (F, H, J) Two-way ANOVA with Šidák correction for multiple comparisons

**Figure 2:** (B) Non-parametric Kruskal-Wallis test of multiple comparisons. (C-E) Two-way ANOVA with Šidák correction for multiple comparisons.

**Figure 3:** (F) Non-parametric Kruskal-Wallis test of multiple comparisons.

**Figure 4:** (D) Non-parametric two-sample Kolmogorov-Smirnov test. (F) Non-parametric Wilcoxon rank sum test.

**Figure 5:** (C) Brown-Forsythe and Welch ANOVA test with Dunnett’s T3 multiple comparisons (% CD25^+^ and % Thy1^+^), non-parametric Kruskal-Wallis test (cell recovery). (D) Unpaired t-test.

**Figure 6:** (B) Brown-Forsythe and Welch ANOVA test with Dunnett’s T3 multiple comparisons. (C) non-parametric Kruskal-Wallis test. (F) Ordinary one-way ANOVA with Dunnett’s T3 multiple comparisons.

**Supplementary Figure 1:** (B, C) unpaired t-test. ns = not significant. (D) Welch’s t-test.

**Supplementary Figure 2:** (A, B) two-way ANOVA with Šidák correction.

**Supplementary Figure 4:** (E) Non-parametric Wilcoxon rank sum test. (G) two-sample Kolmogorov-Smirnov test.

**Supplementary Figure 5:** (A, C) One-way ANOVA (% CD25^+^ and % Thy1^+^), and non-parametric Kruskal-Wallis test (cell recovery). (E) Non-parametric Wilcoxon rank sum test.

**Supplementary Figure 6:** (C) Brown-Forsythe and Welch ANOVA test with Dunnett’s T3 multiple comparisons. (D) non-parametric Kruskal-Wallis test. (E) ordinary one-way ANOVA with Dunnett’s T3 multiple comparisons.

**Supplementary Fig 7:** (A) For every timepoint (days 3, 5, 8, 10), two-way ANOVA with Šidák correction by infection and priming. Significance by infection is shown with * and significance by priming is shown with #. (C) Two-way ANOVA by infection and timepoint, performed either for unprimed conditions or Flt3L-primed conditions. Values that are not significant are omitted.

## Supplementary Tables

**Supplementary Table 1. Unsupervised PCA loading genes and chromatin coordinates.**

PC1 and PC2 loading genes (for bulk RNA-seq) and genome coordinates (for bulk ATAC-seq) along with the individual loading values to compare different populations of *in vivo* and *in vitro* progenitors using PC analysis. Gene expression programs and chromatin accessibility features of expanded HSPCs and their progeny T cell progenitors were compared with previously published *in vivo* progenitor data (*18, 22*).

**Supplementary Table 2. Differential gene expression analysis comparing the transcriptional programs of *in vivo* and *in vitro* progenitors.**

Gene expression features of *in vivo* progenitors from the bone marrow and thymus were compared with those of expanded HSPCs and their progeny pro-T cells generated *in vitro* using OP9-Dll1 co-cultures. Previously published *in vivo* progenitor bulk RNA-seq results (*18, 22*) were utilized for comparisons.

**Supplementary Table 3. Coordinated changes in gene expression and chromatin accessibility programs during the transition from bone marrow progenitor stages to the ETP stage.**

Dynamically regulated genes between expanded HSPCs and their progeny ETPs from OP9-Dll1 co-culture were defined by bulk RNA-seq, and the numbers of different groups of ATAC sites annotated to each gene are shown.

**Supplementary Table 4. Differentially regulated genes between *Trbc1*⁻ and *Trbc1*⁺ Notch-signaled ETPs and association with ATAC changes.**

Gene expression programs of Notch-signaled ETPs were determined from scRNA-seq (normalized counts for *Notch1* ≥ 0.2 & *Hes1* ≥ 0.2 & *Il2ra* < 0.2), and subsets were defined as *Trbc1*⁻ (normalized count <0.2) and *Trbc1*⁺ (normalized count ≥ 0.4) populations. Differentially expressed genes between *Trbc1*⁻ and *Trbc1*⁺ ETPs were shown with the numbers of each group of ATAC peaks.

**Supplementary Table 5. Differentially regulated genes between Lmo2 knockout and control cells in the first five days of T cell development.** Gene expression features from expanded HPSCs co-cultured on OP9-Dll1 for 5 days after either control-infection (DN1) or infection with Lmo2 KO vectors (DN1, DN2). 2 separate gene expression analyses are included: 1) control DN1 vs. Lmo2 KO DN1 and 2) Lmo2 KO DN1 vs. Lmo2 KO DN2.

---

**Supplementary Figure 1.**
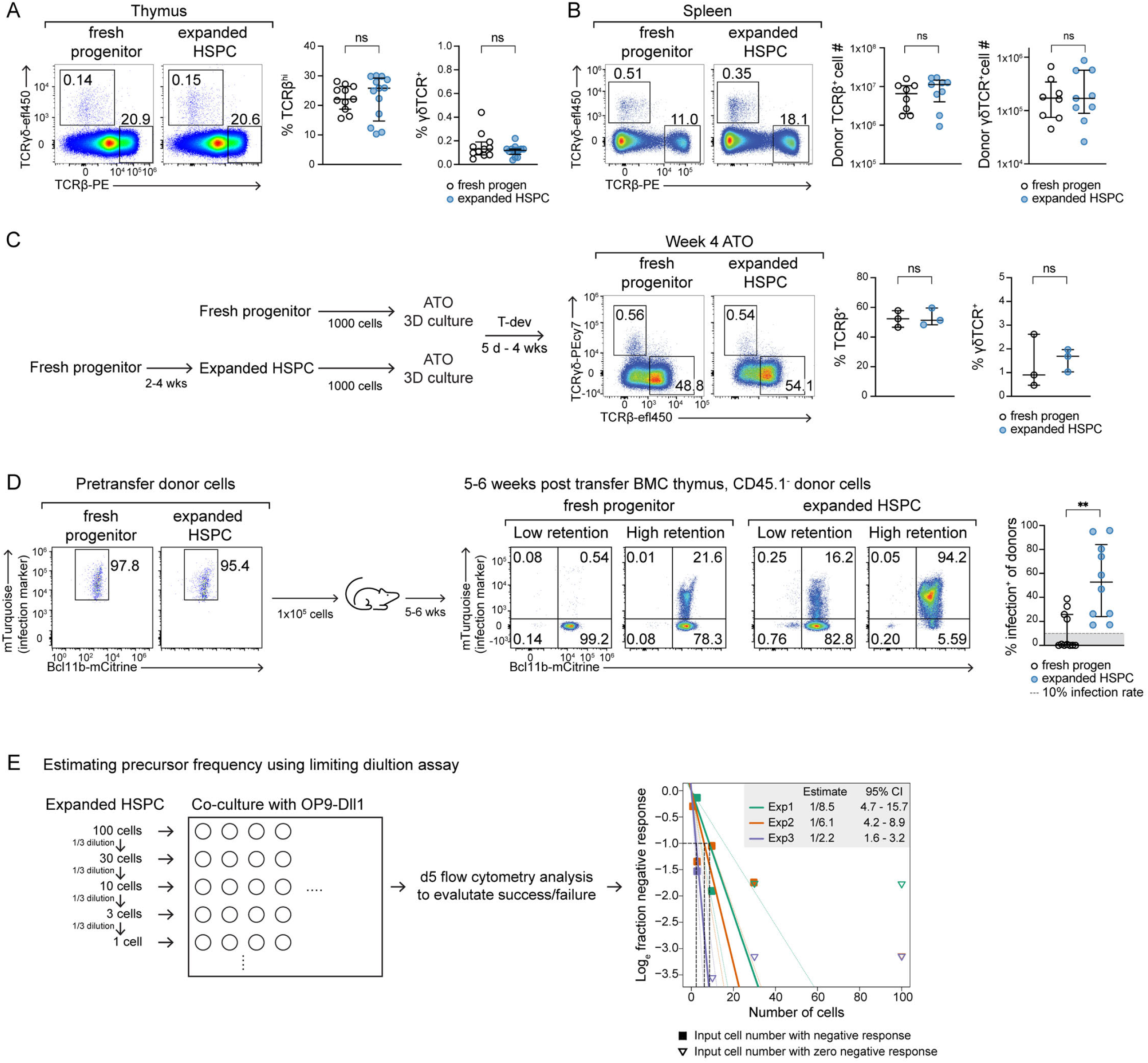
T cell reconstitution from expanded HSPCs *in vivo* and *in vitro* and precursor frequency of expanded HSPCs under T-development conditions. **(A, B)** Bone marrow chimeric mice were reconstituted from freshly isolated progenitors or expanded HSPCs. The expression of TCRβ and TCRγδ of the donor cells in the thymus (A) and the spleen (B) was assessed at 4 weeks post-reconstitution. Representative flow plots show the frequency of TCRβ^hi^ and TCRγδ^+^ cells. Frequencies or cell numbers of indicated populations shown with median and interquartile ranges from 2-3 independent experiments (3-4 mice per group, a total of 11-13 mice for the thymus, and a total of 8-9 mice for the spleen). Statistical test by unpaired t-test. ns = not significant. **(C)** Testing T cell potential of expanded HSPCs in artificial thymic organoids (ATOs). The experimental scheme describes the strategy for ATO culture. Flow cytometry plot represents TCRβ and TCRγδ expression patterns from 3 independent ATO experiments at week 4. Graphs show the percentage of TCRβ^+^ or TCRγδ^+^ cells with median and interquartile range. Each dot represents the average values of 1-5 technical repeat ATOs. Statistical test by unpaired t-test. ns = not significant. **(D)** Retroviral infection marker (mTurquoise) expression patterns in fresh progenitors and expanded HSPCs before and after transfer into busulfan-conditioned recipients are shown. Representative flow cytometry plots display the mTurquoise levels of donor cells from animals with low retention of infection^+^ cells vs. those with high retention of infection^+^ cells in the thymus at 5-6 weeks post-transfer. The graph illustrates the frequency of infection^+^ cells among the donor cells with median values and interquartile ranges. Comparisons by Welch’s t-test. **= *p*-value < 0.01. **(E)** Clonogenic precursor frequency of expanded HSPCs under T-cell conditions was estimated using limiting dilution assay by plating one cell to 100 cells to OP9-Dll1 cells and co-culture for 5 days. The graph shows the estimated precursor frequency (x values corresponding to y = ln(e^-1^), and the 95% confidence interval (CI) ranges for reciprocals of precursor frequencies, from 3 independent limiting dilution assays. The success/failure rate of each cell dose was obtained from 3-24 technical replicate wells. Estimates of precursor frequencies in each experiment, and reciprocals of the 95% confidence limits, were calculated with the Extreme Limiting Dilution Analysis (ELDA) tool (*79*).

**Supplementary Figure 2.**
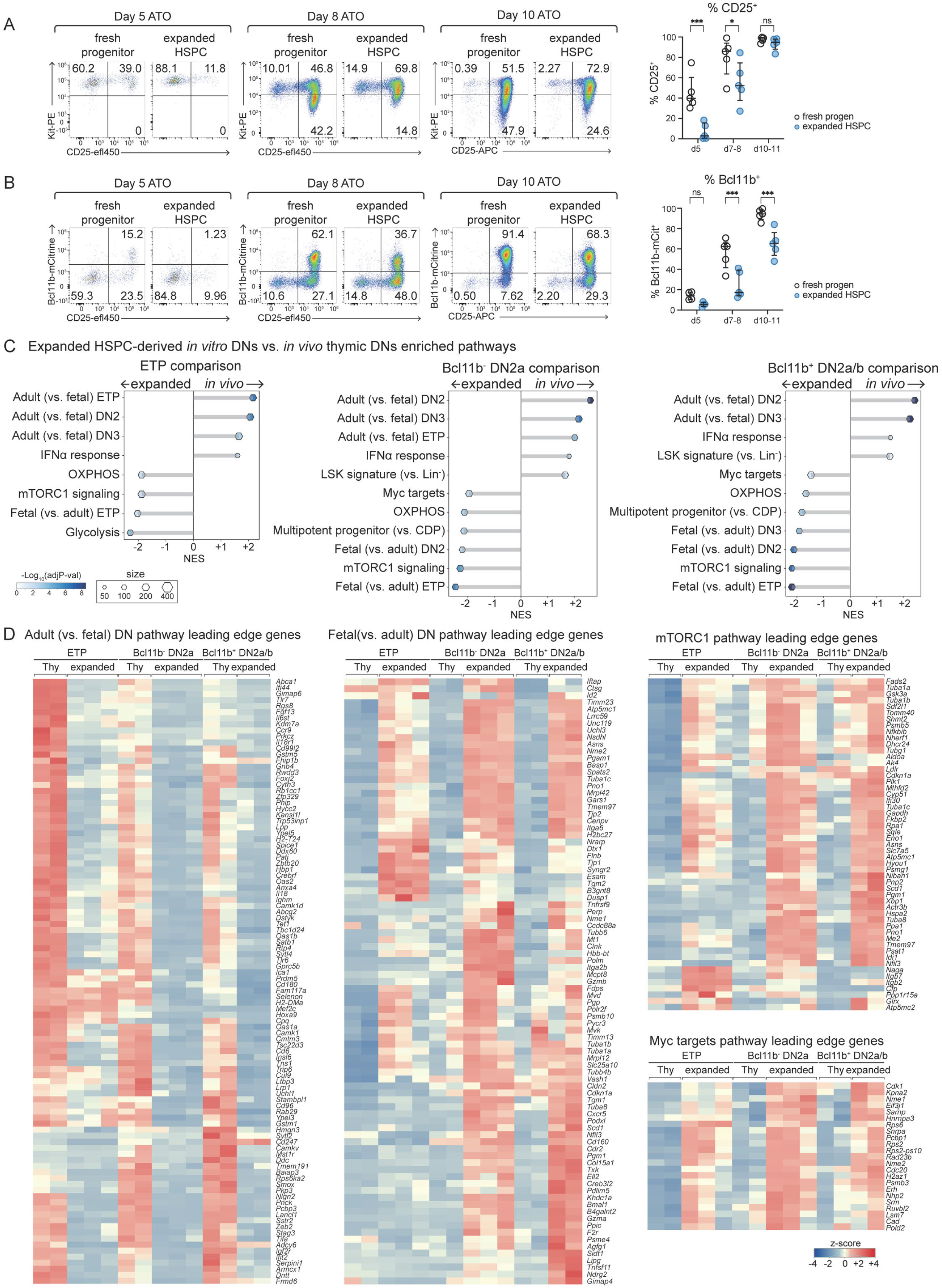
Early T cell development from expanded HSPCs shows a slow progression rate and upregulation of genes associated with active metabolism and fetal T cell development. **(A, B)** ATOs were formed using 1000 cells of fresh progenitor or expanded HSPCs to track early T-developmental progression. The pro-T cell stages were determined by Kit, CD25, and *Bcl11b*-mCitrine reporter expression at days 5, 8, and 10, and the representative flow cytometry plots were displayed. Graphs show the frequency of CD25^+^ cells and Bcl11b-mCitrine^+^ cells at each time point with median values and interquartile ranges. Comparisons by two-way ANOVA with Šidák correction. *=*p-*value < 0.05, *** = *p-*value < 0.001, ns = not significant. **(C)** Gene set enrichment analysis (GSEA) reveals enriched pathways within *in vivo* thymic progenitors vs. expanded HSPC progeny DNs. Normalized enrichment scores (NES) for significantly enriched pathways in ETP, Bcl11b^-^DN2a, and Bcl11b^+^DN2a/b stages are separately plotted. Pathways highly utilized by *in vivo* thymic progenitors have positive NES, and those preferentially employed in expanded HSPC progeny have negative NES. The hexagon size represents the size of the indicated gene set, and the colormap reflects the significance by-log_10_(adj *p-*value). **(D)** The leading-edge genes overlapped with differentially expressed genes (DEG) from *in vivo* thymic DNs vs. *in vitro* expanded HSPC progeny comparisons are shown in the heatmaps. The color map shows the *Z*-score of the TPM value of each gene across the samples.

**Supplementary Figure 3.**
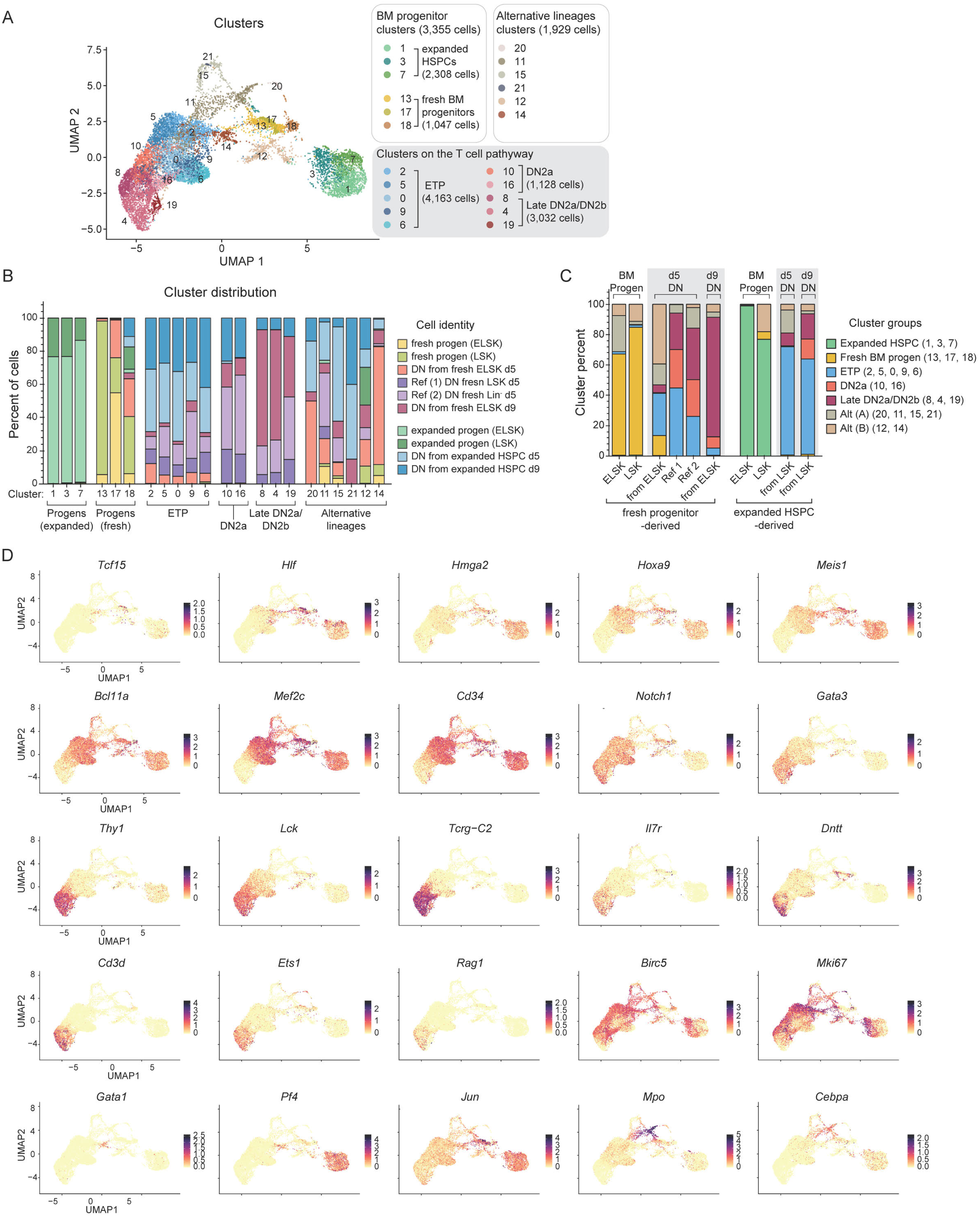
Single-cell transcriptome profiles reveal shared and distinct gene expression programs by pro-T descendants from fresh progenitors and expanded HSPCs. **(A)** Cells from scRNA-seq are displayed in the UMAP1 −2 manifold and colored by Louvain clusters. Cluster groups and the number of cells in each group are noted. **(B)** A stacked bar graph summarizes the membership percentage of cells in each cluster. **(C)** The membership percentage of clusters in each cell type is shown. **(D)** The color intensity in UMAP exhibits expression levels of informative genes representing different progenitor stages, alternative lineage programs, and T cell development.

**Supplementary Figure 4.**
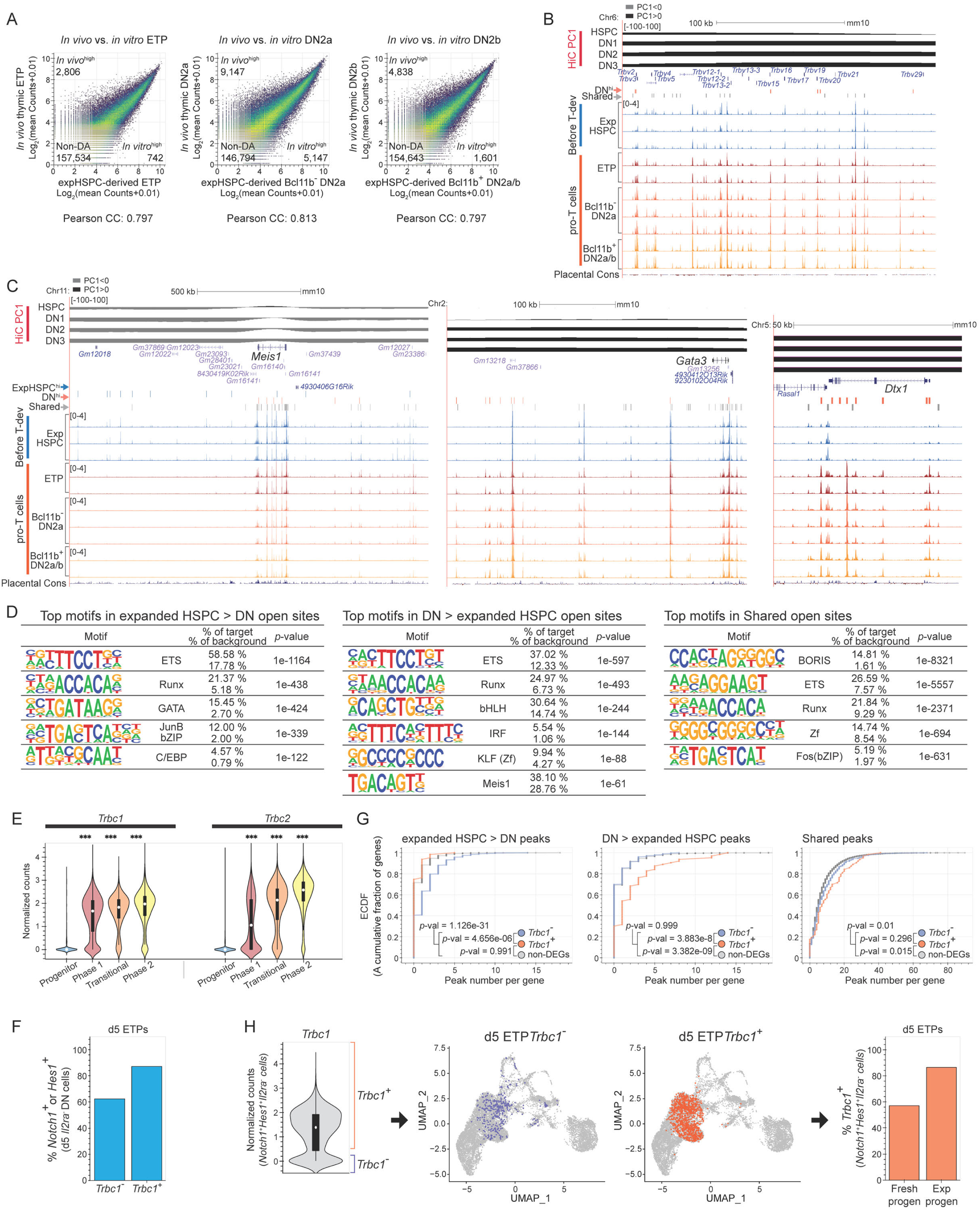
Chromatin accessibility features upon entry to the T cell programs. **(A)** ATAC-signals from *in vivo* thymic DNs and expanded HSPC (expHSPC) progeny DN are compared. Scatter plots show Log_2_-transformed mean counts of each peak from 2-3 experimental repeats for indicated samples. The number of differentially accessible (DA) peaks (absolute fold-change ≥ 2, adjusted *p-*value < 0.01) from *in vivo* (top left) vs. *in vitro* (bottom right), along with the non-DA peak numbers (bottom left), are shown. The linear correlation was measured by the Pearson correlation coefficient (CC). **(B, C)** Representative UCSC genome browser tracks show chromatin accessibility profiles of expanded HSPC (expHSPC) and their *in vitro*-generated pro-T cell descendants (ETP, Bcl11b^-^DN2a, and Bcl11b^+^DN2a/b). The peaks preferentially open in expanded HSPC (expHSPC^hi^, blue), pro-T cells (DN^hi^, orange), and the consistently accessible peaks (shared, grey) are marked above the tracks with thin lines. The 3D chromatin states were determined by previously published HiC (*23*) and represented by PC1 values at the top of the tracks, in which positive values (black) reflect active A compartment and negative values (grey) indicate inactive B compartment. The UCSC Genome Browser’s placental mammal conservation track reflects sequences that are under different evolutionary pressures. The *Trbv* genes (B), *Meis1*, Gata3, and *Dtx1* (C) genomic regions are displayed. **(D)** The top enriched motifs in differentially accessible peaks and shared peaks are demonstrated. **(E)** Normalized counts of *Trbc1* and *Trbc2* transcripts from scRNA-seq are shown. Violin plots show median values (white circle), interquartile ranges (thick bar), and 1.5x interquartile range (thin line). *** = adj *p-*value < 0.001 (compared with progenitor cluster group). **(F)** The bar graph shows the percentage of *Notch1* or *Hes1* expressing (normalized count ≥ 0.2) cells among the *Trbc1-* (*Trbc1* < 0.2, purple) and *Trbc1*^+^ (*Trbc1* ≥ 0.4, orange) day5 ETPs (normalized count for *Il2ra* < 0.2). **(G)** The violin plot demonstrates *Trbc1* expression patterns within d5 Notch-signaled ETPs (normalized count *Notch1* ≥ 0.2 & *Hes1* ≥ 0.2 & *Il2ra* < 0.2). Note the bimodal distribution (left). The *Trbc1^-^* (*Trbc1* < 0.2, purple) and *Trbc1*^+^ (*Trbc1* ≥ 0.4, orange) Notch-signaled d5 ETPs are highlighted in UMAP1-2 (center). The percentage of *Trbc1*^+^ cells of d5 ETPs from fresh progenitor descendants and expanded HSPC descendants are shown (right). **(H)** Empirical cumulative distribution function (ECDF) plots show associations of indicated groups of genes with differentially accessible or constantly open chromatin regions during the development by enumerating the number of peaks near each gene. Distribution comparisons by two-sample Kolmogorov-Smirnov test.

**Supplementary Figure 5.**
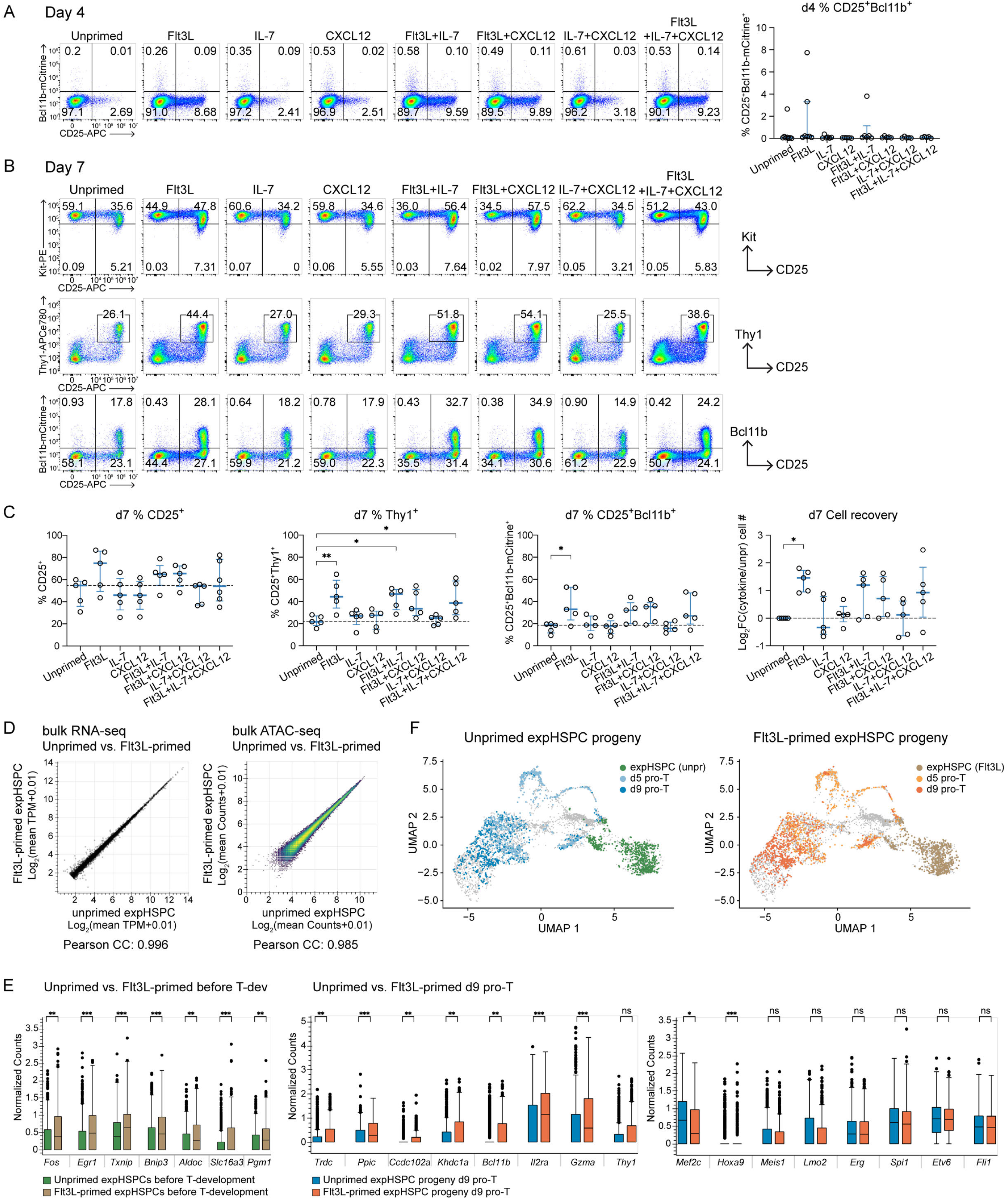
Flt3L treatment in expanded HSPCs assists ETP to DN2 transition without major transcriptional or epigenetic changes. **(A-C)** Expanded HSPCs were treated with different cytokine or chemokine conditions before the *in vitro* OP9-Dll1 co-culture. After 4 days (A) or 7 days (B, C) of culture, T-development stages and cell recoveries were measured. Flow cytometry plots show CD25, *Bcl11b*-mCitrine, Kit, and Thy1 levels, and graphs summarize 5-6 independent experiments with median and interquartile values. Statistical comparisons by one-way ANOVA. *** = *p-*value < 0.001, ** = *p-*value < 0.01, * = *p-*value < 0.05. **(D)** Scatter plots compare bulk RNA-seq (left) and ATAC-seq (right) results of untreated vs. Flt3L-treated expanded HSPCs. Linear relationships were shown by the Pearson correlation coefficient. **(E)** The box plots show differentially expressed genes (DEGs) and non-DEGs of interest in expanded HSPCs in pre-Notch stages (left) and 9 days after T-cell development conditions (right) measured by single-cell RNA-seq. *** = adj *p-*value < 0.001, ** = adj *p-*value < 0.01, * = adj *p-*value < 0.05. DEGs were determined by absolute Log_2_FC > 0.5 by comparing untreated HSPCs with Flt3L-primed HSPCs. **(F)** Single-cell transcriptome results of unprimed and Flt3L-primed expanded HSPCs and their progeny are shown in UMAP.

**Supplementary Figure 6.**
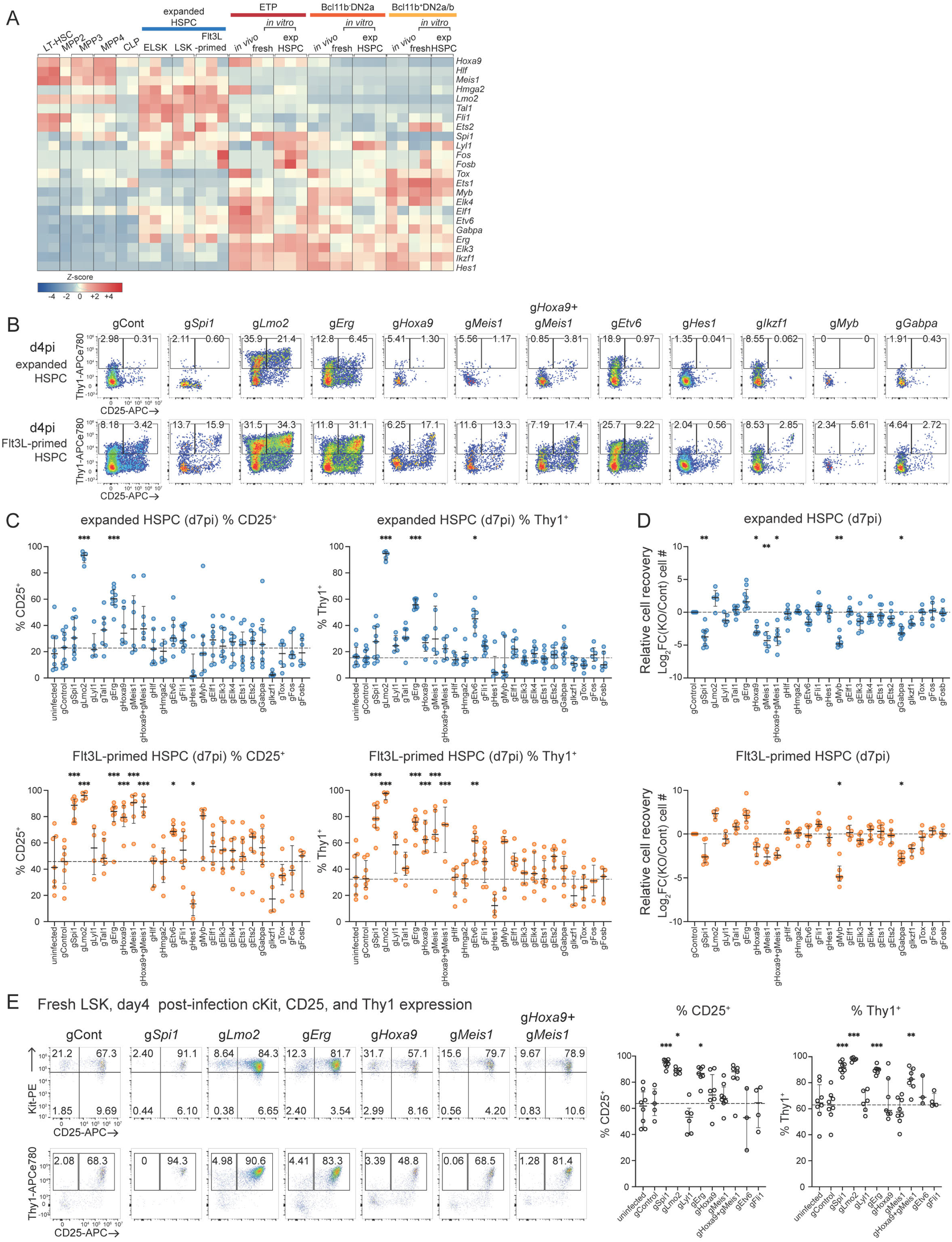
Acute disruption of hematopoietic transcription factors using CRISPR-Cas9 revealed new T-development controllers. **(A)** The expression levels of 23 transcription factors in progenitor stages are illustrated using a heatmap. The *Z*-score of TPM values are plotted. **(B)** CD25 and Thy1 expression levels were measured using flow cytometry after 4 days of indicated TF deletion in unprimed and Flt3L-primed expanded HSPCs. **(C, D)** Graphs show frequencies of Thy1^+^ cells and relative cell recovery after 7 days of OP9-Dll1 co-culture following acute TF deletion. Median values and interquartile ranges were obtained from 5-14 independent experiments. Each data point represents an average of 1-4 technical replicates. Statistical comparisons by Brown-Forsythe and Welch ANOVA test with Dunnett’s T3 multiple comparisons (C) and Kruskal–Wallis test (D). * = *p-*value < 0.05, ** = *p-*value < 0.01, *** = *p-*value < 0.001. **(E)** Acute deletion of indicated TFs in freshly isolated LSKs validates their roles in T-developmental progression after 4 days of gRNA delivery. Representative flow cytometry plots show Kit, CD25, and Thy1 expression levels, and graphs summarize frequencies of CD25^+^ and Thy1^+^ cells. Median values and interquartile ranges from 3-8 independent experiments are shown. Statistical analysis by ordinary one-way ANOVA with Dunnett’s T3 multiple comparisons. * = *p-*value < 0.05, *** = *p-*value < 0.001.

**Supplementary Figure 7.**
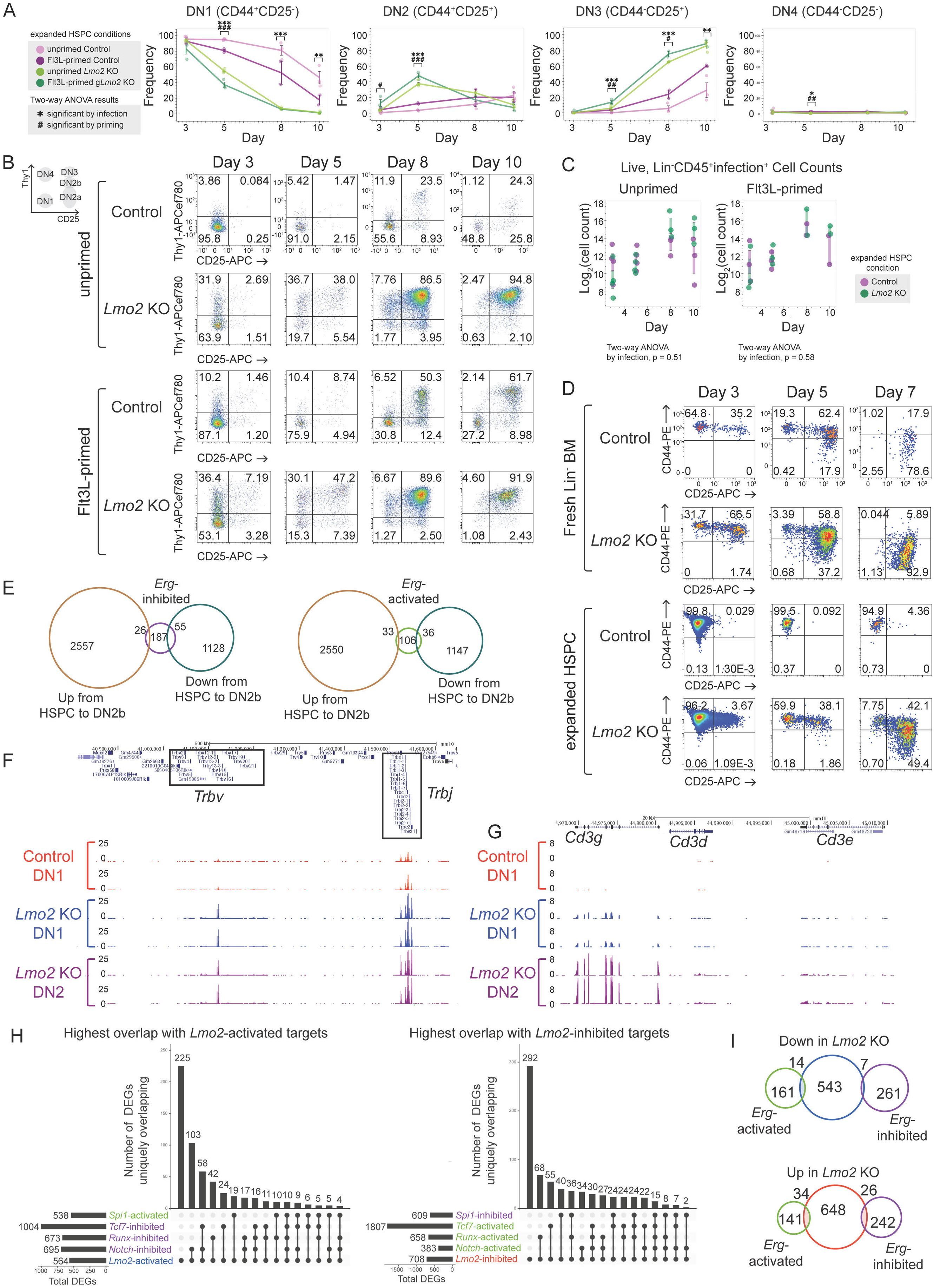
Acute Disruption of *Lmo2* in expanded HSPCs causes a consistent developmental acceleration without survival loss, unique from Erg*-*regulated phenotypes and correlated with other TF activity. **(A)** Different DN frequencies from flow analysis (shown in Fig. 7B) are plotted across all timepoints. 3-4 technical replicates per condition (unprimed and Flt3L-primed, either control or *Lmo2* KO) are shown over 2 independent experiments. Statistical comparisons by two-way ANOVA (considering priming and infection) with Šidák correction for each timepoint. **(B)** Representative flow cytometry plots for experiments shown in Fig. 7A-C display Thy1 and CD25 expression levels. **(C)** Cell counts for the live, singlet, lin^-^, CD45^+^ infection^+^ (mTurq^+^) populations were plotted across all samples. Statistical comparisons by two-way ANOVA (considering infection and timepoint), which show no significant difference by infection. **(D)** Representative flow plots of OP9-Dll1 co-cultures seeded with Lin^-^ bone marrow-derived progenitors or expanded HSCPs, either infected with control or *Lmo2* KO vectors. DN stages shown by CD44 and CD25. **(E)** Venn diagram compares Erg-inhibited and Erg-activated target genes to the reference DEGs from the HSPC to DN2b stage used in Fig. 7E. **(F, G)** UCSC Genome Browser plots show the bulk RNA-seq tracks at the TCRβ-coding locus and the *Cd3* genes for Cont-DN1, *Lmo2*KO-DN1, and *Lmo2*KO-DN2 cells across 2 independent experiments. **(H)** UpSet plots show unique intersections between Lmo2 targets and relevant Notch, Runx, TCF1 (encoded by *Tcf7*), and PU.1 (encoded by *Spi1*) targets. The total number of each TF set is also shown. **(I)** Venn diagram compares Erg-activated and Erg*-*inhibited targets to Lmo2-activated and Lmo2-inhibited target genes.

